# Ecological, angler and spatial heterogeneity drive social and ecological outcomes in an integrated landscape model of freshwater recreational fisheries

**DOI:** 10.1101/227744

**Authors:** S. Matsumura, B. Beardmore, W. Haider, U. Dieckmann, R. Arlinghaus

**Author notes:** ceased after a tragic bike accident before this manuscript was completed (W. Haider). Author for correspondence: Faculty of Applied Biological Sciences, Gifu University, Yanagido 1-1, 501-1193 Japan.

## Abstract

Freshwater recreational fisheries constitute complex adaptive social-ecological systems (SES) where mobile anglers link spatially structured ecosystems. We present a general social-ecological model of a spatial recreational fishery for northern pike (*Esox lucius*) that included an empirically measured mechanistic utility model driving angler behaviors. We studied emergent properties at the macro-scale (e.g., region) as a result of local-scale fish-angler interactions, while systematically examining key heterogeneities (at the angler and ecosystem level) and sources of uncertainty. We offer three key insights. First, the angler population size and the resulting latent reginal angling effort exerts a much greater impact on the overall regional-level overfishing outcome than any residential pattern (urban or rural), while the residential patterns strongly affects the location of local overfishing pockets. Second, simplifying a heterogeneous angler population to a homogenous one representing the preference and behaviours of an average angler risks severely underestimating landscape-level effort and regional overfishing. Third, we did not find that ecologically more productive lakes were more systematically overexploited than lower-productive lakes. We conclude that understanding regional-level outcomes depends on considering four key ingredients: regional angler population size, the angler population composition, the specific residential pattern in place and spatial ecological variation. Simplification of any of these may obscure important dynamics and render the system prone to collapse.

## Introduction

Recreational fishers are the dominant users of wild living fish stocks in most inland fisheries and many coastal ones in the industrialized world (Arlinghaus et al., 2015). Recreational fisheries constitute complex adaptive social-ecological systems (SESs) (Hunt et al., 2013; Ziegler et al., 2017), which are characterized by three key features (Arlinghaus et al., 2017): individual and spatial heterogeneity, hierarchical organization across scales (from micro to macro levels) and the presence of non-linearities leading to the potential for regime shifts. Outcomes in complex adaptive SESs at macro-scales (e.g., regionally, nationally or globally) are an emergent, self-organized property of local-level interactions among humans and ecosystems (Levin et al., 2013). For example, in open-access freshwater recreational fisheries local, micro-level interactions of anglers and selected lakes or river sections give rise to a spatial spread of angling effort on the macro-level as anglers select sites that promise high utility. Alternatively framed, the dynamic site choice behaviour of anglers at equilibrium produces regional-level outcomes at the macro-scale, such as degree of overfishing, spread of non-natives fishes and social well-being or conflict (Arlinghaus et al., 2017). If we are to advance our understanding of recreational fisheries as complex adaptive SESs, a focus on the macro-scale outcomes and how they mechanistically result (i.e., emerge) from a range of micro-scale feedbacks among anglers and fish stocks/ecosystems is needed (Arlinghaus et al., 2017). This is particularly the case in freshwater recreational fisheries where there is an exceedingly large individual (i.e., angler-level) and spatial heterogeneity (i.e., among lake variation in ecological quality) and where cross-scale feedbacks among social and environmental subsystems are commonly observed (Arlinghaus et al., 2017; Mee et al., 2016; Wilson et al., 2016; Ziegler et al., 2017).

One characteristic, yet managerially largely overlooked feature of most freshwater recreational fisheries is their spatial structure, both in terms of spatial variation in productivity of different ecosystems (Kaufmann et al., 2009; Lester et al., 2003; Post et al., 2008; Shuter et al., 1998) as well as spatial variation in residential patterns of the human forager in terms of where anglers live relative to the available resource patches (lakes, river section) they seek. Broadly speaking, a water-rich freshwater fisheries landscape can be exploited by human foragers living a small number of large metropolitan areas (e.g., Post et al., 2008) or human foragers may reside in rural contexts in a multitude of individual villages and towns spread in the landscape. The residential structure affects travel costs, which is a key component of angler utility and hence site choice (Hunt 2005; Post et al., 2008). Therefore, the fishing pressure on any given locality will be a function of where the forager population is geographically located relative to the locality, but no systematic research is available on this topic.

Overall, few studies on the landscape dynamics of freshwater recreational fisheries exist, most of which are from North America (Askey et al., 2013; Carpenter & Brock, 2004; Hunt et al., 2011; de Kerkhove et al., 2015; Mee et al., 2016; Post & Parkinson, 2012; Post et al., 2008; Shuter et al., 1998; Wilson et al., 2016; Ziegler et al., 2017). Most of these studies were focused on how overfishing and other regional outcomes related to an urban residential pattern of the human forager population (e.g., Carpenter & Brock, 2004; Hunt et al., 2011; Post et al., 2008), modelled one specific landscape characterized by a unique management approach (e.g., stocking-based rainbow trout, *Oncorhynchus mykiss,* fishery in British Columbia near Vancouver, Post et al., 2008) and omitted addressing systematic effects of heterogeneity within the forager (angler) population by focusing on angling effort as an aggregate outcome (Camp et al., 2015; Post et al., 2008). This is a relevant shortcoming because angler diversity in preferences and behaviour is likely to strongly affect feedbacks among social and ecological subsystems (Johnston et al., 2010) and thereby dictate regional-level outcomes (e.g., where overfished stocks are expected to happen, Hunt et al., 2011).

To further our understanding about which outcomes to expect from the localized interaction of fish and anglers at the landscape scale, the construction of process-based simulation models carrying sufficient mechanistic detail about the main driving mechanisms (e.g., compensatory reserve of fishes varying in productivity across lakes or site choice process exhibited by heterogeneous anglers) is needed (Fenichel et al., 2013a; Schlüter et al., 2012). Process-based modelling approaches seem warranted because the complex adaptive system of recreational fisheries is characterized by many non-linear feedbacks whose joint effects are difficult to be predicted beyond the sphere of observed parameters in correlation-based models (Arlinghaus et al., 2017; Fenichel et al., 2013a; Hunt et al., 2011; Schlüter et al., 2012;). One key ingredient to include in models of the SES of recreational fisheries is a mechanistic model of angler behaviour (Abbott & Fenichel, 2013; Allen et al., 2013; Fenichel et al., 2013a; Johnston et al., 2015). Explicitly representing the mechanisms of site choice by anglers and how site-choice behaviour is affected by both catch and non-catch related experience preferences can lead to strongly differing predictions about the distribution of foragers and ultimately regional-level outcomes compared to models where the behaviour of anglers is simplified to those determinants that would drive natural foragers, e.g., expected catch rates (Hunt et al., 2011; Johnston et al., 2010; Matsumura et al., 2010).

We aimed at studying regional-level overfishing patterns and social outcomes in a rich class of recreational fisheries landscapes that varied in the geographical distribution of the human forager population, using a general social-ecological model that involved an empirically estimated mechanistic model of site choice of anglers, while accounting for ecological variation among lakes and angler heterogeneity in preferences and behaviour. We wanted to go beyond existing landscape investigations that were usually tailored towards one specific residential pattern and geography and thereby provide general insights into which spatial patterns of effort, yield, angler well-being and overfishing to expect in varying residential scenarios, for varying angler population sizes and for anglers and lakes varying in key features (preferences or productivities, respectively). We sought answers to three key questions:

1. Which systematic impact on regional-level outcomes in an open-access freshwater recreational fishery can be expected from variation in residential patterns of the human forager population ranging from urban to rural?
2. Which systematic effects on regional-level outcomes can be expected to arise from heterogeneity in angler preferences and behaviour?
3. Which systematic effects on regional-level outcomes can be expected to arise from among-lake ecological heterogeneity in productivity and carrying capacity?

Related to these three objectives, we hypothesized (H1) that a rural residential pattern will even out landscape-level overfishing and render the placement of overfished stocks less concentrated around urbanities (the latter was usually reported from urban fisheries landscapes, Hunt et al., 2011; Post et al., 2008), (H2) that angler heterogeneity will aggravate regional overfishing by spreading effort in space to also remote fisheries (Ward et al., 2013b), and (H3) that we will continue to find little evidence for more productive fisheries being systematically overexploited by anglers that follow a multi-dimensional utility function when searching for fishing sites in space (Hunt et al., 2011).

The rationale for the third and last hypothesis is that anglers are known to choose lakes following a multi-dimensional utility function where various non-catch dimensions of the angling experience (e.g., social aspects, distance, costs, harvest regulations) affect the expected utility of a site or ecosystem to anglers in addition to those dimensions that are strongly about catch expectations (e.g., catch rate, size of the fish that are captured) (Arlinghaus et al., 2014; Cole & Ward, 1994; Hunt 2005; Johnson & Carpenter, 1994; McFadden, 1973). Moreover, anglers are known to be highly heterogeneous in their preferences and behaviours (Anderson, 1993; Beardmore et al., 2011; Cole & Ward, 1994; Dorow et al., 2010; Fenichel & Abott, 2014; Johnston et al., 2010; Wilde & Ditton, 1994), which will strongly affect where in space a particular angler type will be attracted to (Hunt et al., 2011; Ward et al., 2013a, b). If anglers, however, would be mainly attracted to a given fishery by the expected catch rates with only minor importance attached to other attributes of the lake and the fishing experience in general (e.g., distance, crowding), the classic ideal free distribution framework (Fretwell & Lucas 1970) from behavioural ecology would allow the clear-cut prediction that lakes offering higher catch rates (i.e., more productive fisheries) should be systematically overexploited (Parkinson et al., 2004). Our own pervious work, however, has revealed that such expectations are not warranted (Hunt et al., 2011). Instead, deviations from a catch-based ideal free distribution (where at equilibrium all lakes supposedly offer a regional average catch quality, Mee et al., 2016) should be the norm 1) when angler’s site choice is sub-optimal (by choosing the lake with the highest expected utility probabilistically rather than deterministically), and 2) when multiple attributes in addition to catch affect site choice. Both dimensions – suboptimal lake choice and multiple non-catch attributes providing utility – should foster a dynamic equilibrium that maintains between-lake variation in catch rates and other measures of catch qualities (Hunt et al., 2011; Matsumura et al., 2010), but this predictions remains to be fully explored in the present paper.

We designed our work to provide a comprehensive examination of the systematic impacts of spatial and angler heterogeneity assuming a mechanistic model of angler behaviour following utility theory. Our research is meant to constitute a strategic modelling experiment (as opposed to a tactical modelling approach that looks for insights in relation to a very specific fisheries landscape) about social and ecological regional-level outcomes to be expected when anglers interact in a localized fashion with spatially structured lakes. The behavioural model of angler site choice we use was informed by empirical data from stated behaviour of anglers in Germany (Beardmore et al., 2013), and the fish biological component was calibrated to empirical data of the northern pike *(Esox lucius).* We assume anglers are human foragers, who seek fitness in utility units. We choose pike as the target species due to its circumpolar distribution in most lakes of North America and Eurasia and because pike is a heavy sought species by many anglers across its native range (Arlinghaus & Mehner, 2005; Crane et al., 2015). Despite this calibration, our model provides generic insights into outcomes to be expected from individual and spatial heterogeneity in a coupled SES of recreational fisheries. Results of our work are to be seen as hypotheses to be explored in specific fisheries and as explanation for empirical findings reported elsewhere (e.g., Mee et al., 2016). We hope to provide an innovation over existing SES models of recreational fisheries by presenting several outcomes jointly, related to regional-level ecological objectives (e.g., regional overfishing), regional-level economic objectives (e.g., regional angler welfare) and more traditional fisheries objectives (e.g., average catch rates and effort distribution). Thereby, our work contributes to the importance of being explicit about management objectives in assessing regional-level outcomes of fish-angler interactions (Fenichel et al., 2013b). Finally, our work also offers some strategic management implications into expected ways how traditional management tools designed to affect either people (through harvest regulations) or fish stocks (through activities such as habitat enhancement or stocking) may play out when anglers and fish stocks thriving in spatially and ecologically varying ecosystems are linked through site choice behaviour of a heterogeneous angler population in freshwater landscapes.

### The model Spatial structure

We designed a freshwater fisheries landscape *in silico,* constructing a two-dimensional square lattice of 11 × 11 (=121) lakes, each of a small size of 10 ha. The size was chosen so that angler crowding would be present at high-use fisheries, which reduces attractiveness of a lake (Arlinghaus et al., 2014; Hunt 2005). The distance to a closest neighbouring lake was assumed to be 15 km. We present two extreme residential patterns - uniform (“Rural”) and concentrated (“Urban”). In the rural case, anglers were assumed to live in towns (of identical population sizes) adjacent to lakes across the landscape. In the concentrated urban case, all the anglers were assumed to live in a large city located nearby the central lake of the lattice. We also examined intermediate cases as larger cities (harbouring anglers) scattered through the landscape. As these intermediate cases were found to be always intermediate to the rural and the urban cases, we decided to not present the data in this paper to simplify the presentation. Following the pioneering landscape studies of Carpenter and Brock (2004) and Hunt et al. (2011) and arguments expressed elsewhere (Fenichel et al., 2013a; Johnston et al., 2010), anglers were assumed to move between spatially segregated and ecologically independent lakes according to the (multidimensional) utility each lake provides (for details, see further below). In behavioural ecological terms, the human forager was assumed to select a lake according to the “fitness” offered by a patch (lake) (as assumed in the ideal free distribution theory, Fretwell & Lucas, 1970), with fitness being defined as utility units to anglers rather than prey intake rate as would be the case in natural forager.

### Fish population dynamics

To represent fish populations striving in each of the ecologically unconnected 121 lakes, we used an age-structured model with multiple density-dependent population regulation processes affecting survival and growth and size-dependent survival and fecundity, parameterized with empirical data for pike (Tables 1, 2, Fig. 1). The model is fully presented elsewhere (Arlinghaus et al., 2009, 2010; Matsumura et al., 2011). Briefly, pike growth was modelled with a bi-phasic growth model (Lester et al., 2004; equation 1 in Table 1), where juvenile growth rate is a function of biomass density following empirical data from Windermere (UK). Changes in juvenile growth affect post-maturation growth and the final length that can be attained (Lester et al., 2004). Changes in the biomass density not only affect body length but also fecundity in a density-dependent fashion as reported for pike (Craig & Kipling, 1983).

**Table 1.**
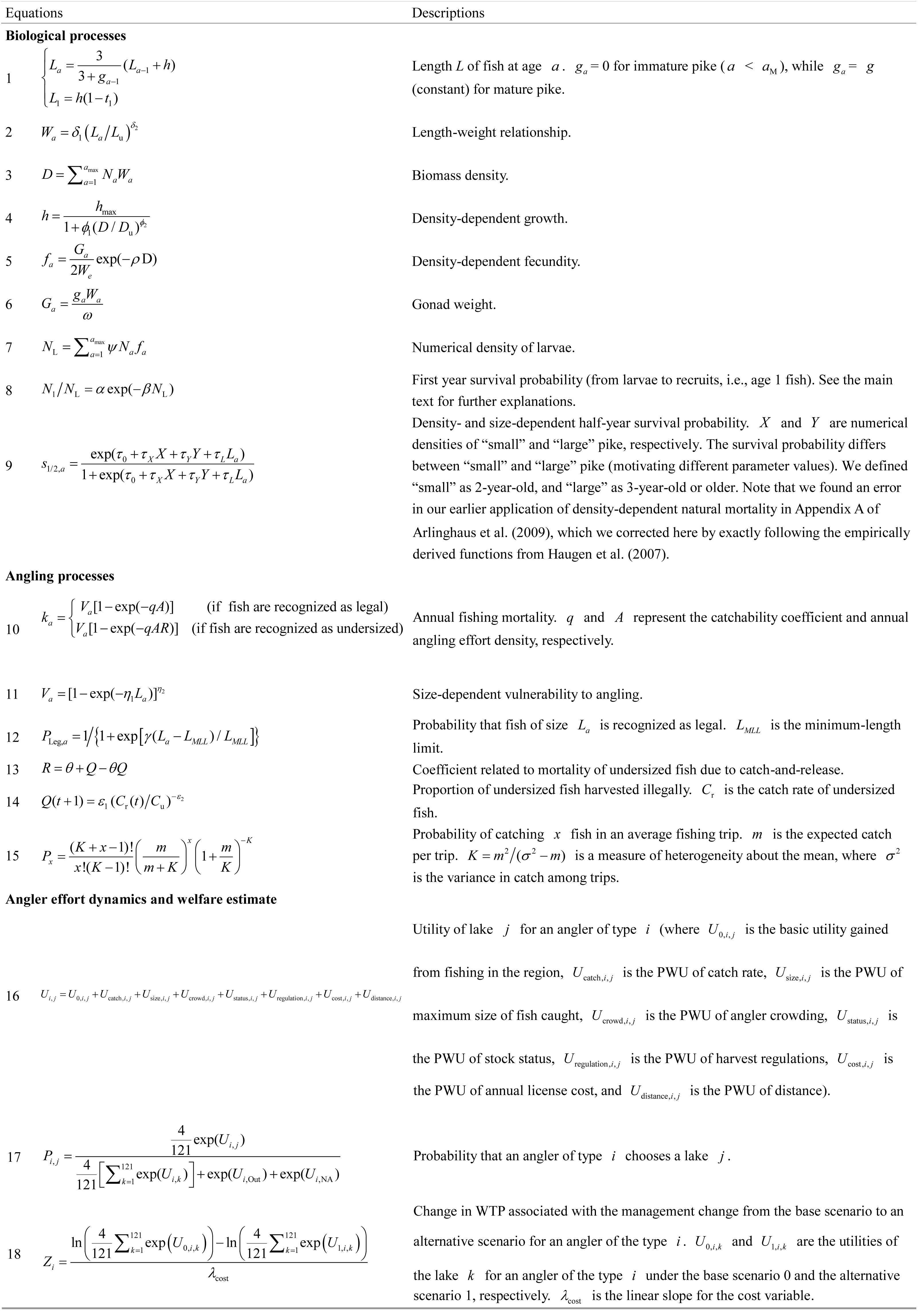
Equations of biological and angling processes of the pike population model (from Arlinghaus et al., 2009). Parameter values and their sources are listed in Table 2.

**Table 2.**
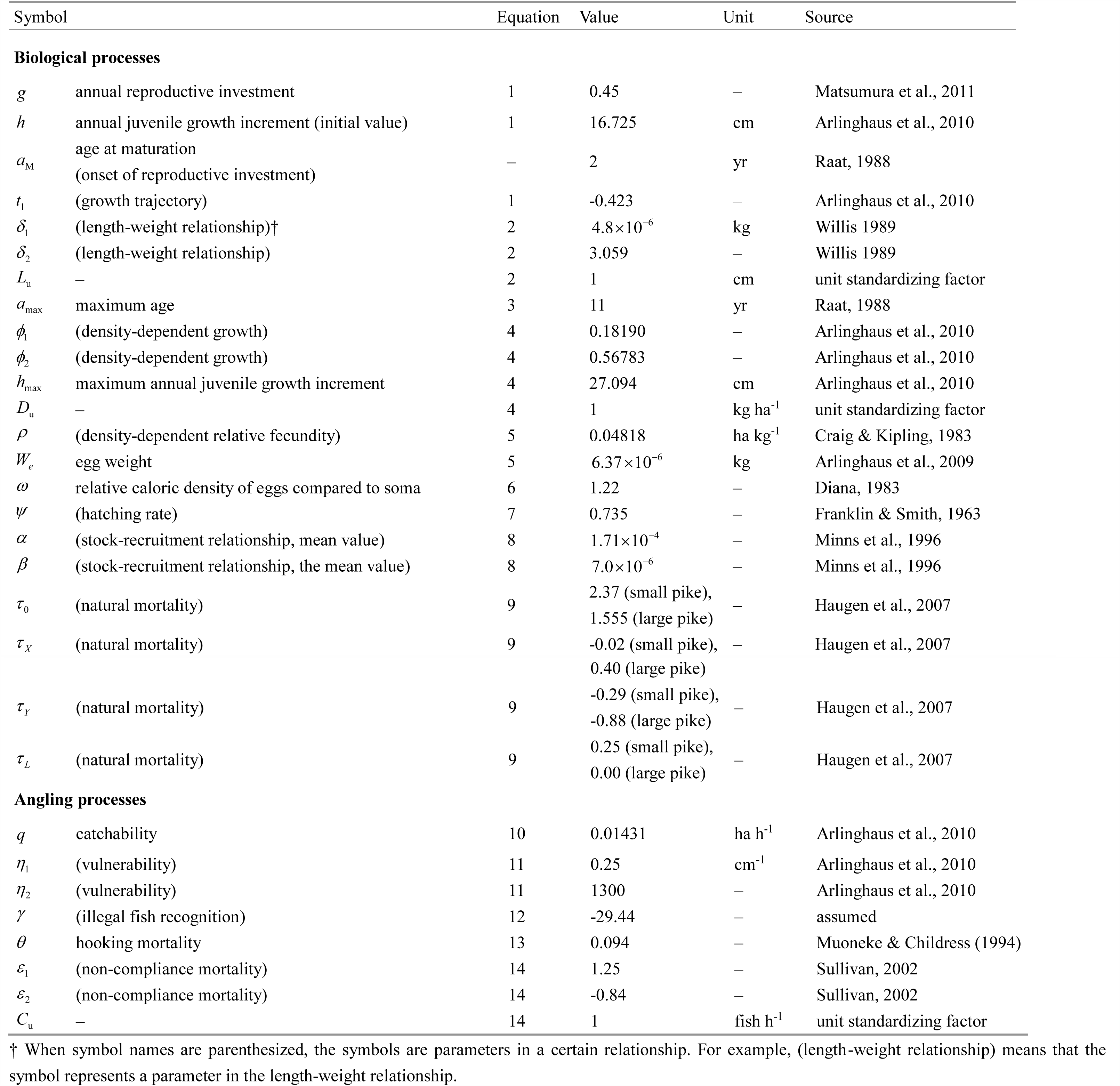
Parameters and parameter values of biological and angling processes.

**Figure 1.**
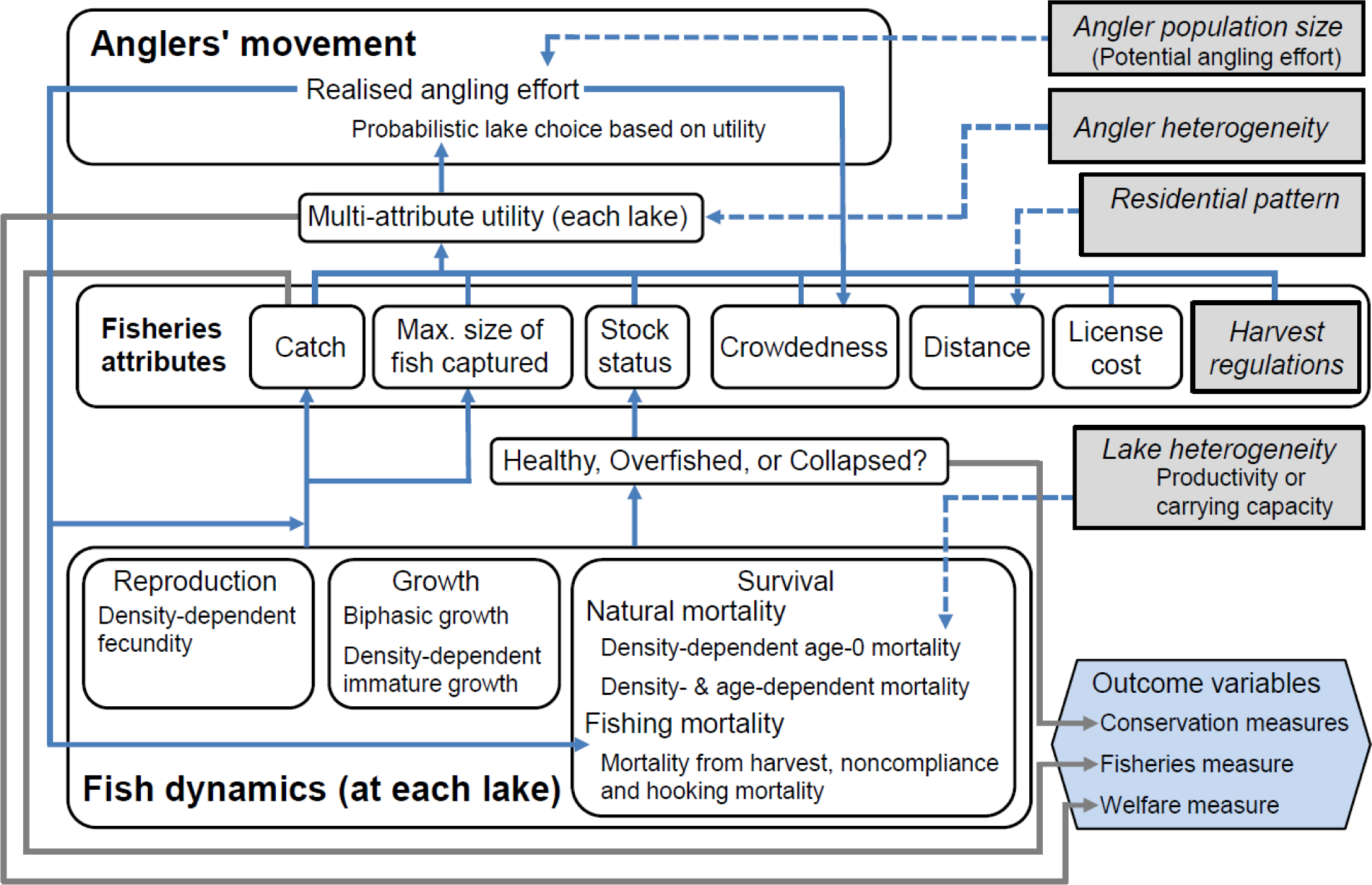
Schematic overview of the model.

The first year survival was modelled using a stock-recruitment relationship assuming Ricker stock-recruitment typical for cannibalistic species such as pike (Edline et al., 2007) of the form

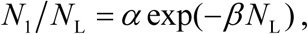

where *N*_1_ and *N*_L_ represent the density of age-1 fish and hatched larvae, respectively; *α* defines the maximum survival rate from spawning to recruitment (i.e., age-1) at low spawner density, and *β* describes the strength of density-dependent interactions influencing the cohort’s survival (Walters & Martell, 2004). Alternatively termed, *β* is the rate of decrease of recruits/spawner as spawner density increases. Both *α* and *β* determine the intrinsic quality of the pike stock, but only *α* strongly affects the slope of the stock-recruitment curve near the origin and thus the per capita number of recruits at low population density (Walters & Martell, 2004). By contrast, *β* determines the maximum recruitment and has little effects on the slope near the origin. As most pike stocks were exploited in our model and hence had lower (exploited) spawning stock biomasses than the virgin population sizes, *α* determines the resiliency of the stock to harvest at low stock sizes and thus the population’s productivity in the exploited state. By contrast, *β* mainly shapes the carrying capacity of a lake for recruits and not the per capita production of recruits at low population sizes. Consequently, we used among-lake variation in *α* to represent variation in productivity of pike stocks, and variation in *β* to represent variation in carrying capacity among lakes. Parameter values of the stock-recruitment function (i.e., the mean values of *α* and *β*) were determined to approximate an empirical relationship reported by Minns et al. (1996) for pike (Table 2).

The pike populations in the 121 lakes differed either in productivity, represented by the parameter *α* (which as above mainly governs the slope of the stock-recruitment relationship at low stock sizes), or in the stock’s carrying capacity, represented by the parameter *β* (which as above governs the maximum number of recruits). The variation of the two parameters represented lake heterogeneity in pike population biology and was assumed to follow a lognormal distribution around a mean. The spatial distribution of lakes was assumed independent of the lake’s biological properties (productivity or carrying capacity), i.e. there was no correlation in the pike stock’s biological properties in neighbouring lakes.

Natural survival after year one was modelled using a size- and density-dependent empirical relationship published for Windermere pike by Haugen et al. (2007) (equation 9 in Table 1). Fishing mortality was modelled with a standard catch equation (equation 10 in Table 1) where catch is determined by effort, abundance and the (constant) catchability coefficient typical for pike (Arlinghaus et al., 2009). Captured fishes were taken home unless protected by regulations, in which case some level of mortality happened due to catch-and-release mortality and non-compliance mortality with regulations following empirical findings for freshwater predatory fish captured by anglers (Muoneke & Childress, 1994; Sullivan, 2002) (equations 13 and 14 in Table 1). Further details on the model can be found in the publications mentioned above as well as Table 1.

### Mechanistic model of site choice by anglers

We followed economic utility theory when designing a model to represent a probabilistic-based site choice behaviour by anglers (Fenichel et al., 2013a; Hunt et al., 2011; Johnston et al., 2010). We choose the most general (i.e., species independent) multi-attribute utility model published so far on recreational anglers when they are confronted with the choice of choosing lakes in space as a function of travel distance and other utility-determining attributes of the fishing experiences, such as expected catch rate, expected size, regulations, crowding and biological status of the fish stock (Beardmore et al., 2013; Johnston et al., 2015). To explore the heterogeneity among anglers, the raw data of the choice experiment by Beardmore et al. (2013) were analysed with a latent class choice model as well as in an aggregate fashion to come up with the average angler in the population (see supporting information). Latent class models statistically determine groups that are maximally different in their preferences (Swait, 1994). We found that a 4-class model explained the data statistically well, which divided anglers into four types in terms of maximal variation in site choice preferences; these anglers were classified in three angler types varying by degree of recreational specialization (from casual to committed, see Johnston et al., 2010 for a summary) in the study region and one highly specialized angler that had a preferences for fishing intensively beyond the study region (see supporting information for details). To study how this heterogeneity of anglers affected our model outcomes, we also studied the exploitation patterns of homogeneous anglers (1-class model) where all the anglers are assumed to be equal in their preferences. Including heterogeneity directly followed the framework of Johnston et al. (2010, 2013, 2015) assuming that anglers vary in importance (the so-called part-worth utility, PWU, estimated from the random utility model, see supporting information for details) attached to specific attributes of the fishing experience and hence behave differently as the fishing environment changes. Estimated parameter values of the 4-class (heterogeneous anglers) and 1-class (homogeneous anglers) models are shown in Table 3.

**Table 3.** Parameter estimates of the latent class preference model for anglers estimated from the choice data presented in Beardmore et al., (2013). Values which are not underlined represent the slope of the PWU (part worth utility) functions, while underlined values represent constants. SD = standard deviation, refers to the standard deviation of data collected from diaries in the study region that were used when standardizing the attribute levels in the original choice experiment. For details on the interpretation of the four angler types, see supporting information.

|  |  |  | 4-class model |  |  |  | 1-class model |
| --- | --- | --- | --- | --- | --- | --- | --- |
| | | | $\lambda$ | | | | $\lambda$ |
| Attribute | Type | Unit | Type 1 | Type 2 | Type 3 | Type 4 | (%) |
|  |  |  | 51.4 | 22.9 | 16.6 | 9.1 | 100.0 |
| Intercept | Nominal | Fish in the region | <u>0.9149</u> | <u>0.1883</u> | <u>-0.5628</u> | <u>-0.4228</u> | <u>0.0900</u> |
|  |  | Fish outside region | <u>-0.2386</u> | <u>-1.0336</u> | <u>-1.1146</u> | <u>1.1449</u> | <u>-0.4637</u> |
|  |  | Not fish at all | <u>-0.6763</u> | <u>0.8453</u> | <u>1.6774</u> | <u>-0.7221</u> | <u>0.3737</u> |
| Catch rate | Numeric | SD from mean | 0.1760 | 0.2337 | 0.4546 | 0.2776 | 0.2040 |
| | | Parameter 1 ($\gamma_1$)<sup>† 1</sup> | 4.1162 | 2.8245 | 1.2646 | 2.1866 | 3.3395 |
| | | Parameter 2 ($\gamma_2$)<sup>† 1</sup> | 2.6299 | 2.7162 | 7.7092 | 3.1479 | 2.6687 |
| | | Parameter 3 ($\gamma_3$)<sup>† 1</sup> | -0.6834 | -0.9623 | -8.6992 | -1.4658 | -0.8141 |
| Maximum size<sup>† 2</sup> | Numeric | SD from mean | 0.1458 | 0.1482 | 0.1699 | 0.1997 | 0.1324 |
| Anglers seen<sup>† 3</sup> | Numeric | SD from mean | -0.0615 | -0.1216 | -0.0982 | -0.1188 | -0.0929 |
| Stock status<sup>† 4</sup> | Nominal | No information | <u>0.0821</u> | <u>0.0940</u> | <u>-0.0225</u> | <u>0.1580</u> | <u>0.0674</u> |
|  |  | Good | <u>0.3087</u> | <u>0.3614</u> | <u>0.5258</u> | <u>0.2674</u> | <u>0.3203</u> |
|  |  | Lightly overfished | <u>-0.0609</u> | <u>-0.0452</u> | <u>0.1618</u> | <u>-0.0961</u> | <u>-0.0338</u> |
|  |  | Overfished | <u>-0.3299</u> | <u>-0.4102</u> | <u>-0.6651</u> | <u>-0.3293</u> | <u>-0.3539</u> |
| Regulations<sup>† 5</sup> | Nominal | None | <u>-0.1660</u> | <u>-0.0544</u> | <u>-0.0675</u> | <u>-0.5259</u> | <u>-0.1307</u> |
|  |  | Light | <u>0.1462</u> | <u>0.0570</u> | <u>0.0657</u> | <u>0.4538</u> | <u>0.1604</u> |
|  |  | Medium | <u>0.0937</u> | <u>0.1020</u> | <u>0.2497</u> | <u>0.1268</u> | <u>0.0799</u> |
|  |  | Strict | <u>-0.0738</u> | <u>-0.1047</u> | <u>-0.2479</u> | <u>-0.0547</u> | <u>-0.1097</u> |
| Licence cost | Numeric | 10 € increment | -0.0212 | -0.0981 | -0.1501 | -0.1251 | -0.0621 |
| Distance | Numeric | 20 km increment | -0.0558 | -0.5212 | -1.6055 | -0.2627 | -0.1293 |

†1: Parameters for the PWU non-linear function, see supporting information for details. †2: Maximum size of fish captured is defined as the 95th percentile of the size distribution of fish caught annually. †3: All anglers at a particular lake are assumed to see each other because of the small size of lakes (10 ha). †4: In the present study, the level “none” (no information present) was chosen (see text for details). †5: The level "Medium" and "None" correspond to the "traditional harvest regulation" scenario (a minimum-length limit of 50 cm and a daily bag limit of 3 pike per angler day) and the "no regulation" scenario, respectively.

In our simulations, anglers were assumed to choose a fishing site (i.e., a lake) offering maximum utility compared to all other utilities offered by all other lakes and to move to the lake with the highest utility probabilistically (equations 16 and 17 in Table 1). Note that although this model assumed utility maximization and perfect knowledge of the utility offered by all lakes, the actual choice was not deterministic but probabilistic (i.e., suboptimal) (equation 17 in Table 1), similar to Matsumura et al. (2010) and Hunt et al. (2011). This agrees with the assumption of bounded rationality common to humans. The weighing factor 4/121 in the equation reflected the fact that survey respondents in the stated choice experiment by Beardmore et al. (2013) had four alternative lakes in the region in addition to the options for fishing outside the region and no fishing (see supporting information), while our virtual anglers had a choice of 121 lakes in their landscape.

In the simulations, anglers were assumed to have perfect information about the average fish number to be expected from each lake, the maximum size of fish to be expected at each lake, and the number of anglers seen at each lakes using information from the preceding year. This might be considered unrealistic, but novel communication means permit spread of information about expected catch rates and other lake attributes quickly. However, we did not consider knowledge about stock status to affect angler choice because it is unrealistic that managers can derive this information every year; we thus kept the attribute value at “no knowledge” in all simulations (Table 3). The maximum size of fish captured at each lake was defined as the 95th percentile of the size distribution of fish caught at the lake during the preceding year. All anglers at a particular lake were assumed to see each other because of the small size of lakes (10 ha). The annual licence cost for angling in the region was fixed at 100 €, which represents a typical value for licence money in Germany (Arlinghaus et al., 2015).

### Regional outcome metrics

We kept track of a range of social, economic and ecological outcome metrics at the regional level used to assess the emergent properties of fish-angler interactions at the landscape levels.

In terms of social and economic metrics, the choice experiment included two dimensions of monetary costs that can be used to quantify the (realized) utility of fishing offered at equilibrium. One was related to travel distance and one related to the direct inclusion of a monetary cost variable (i.e., annual license cost in Euro). The coefficient estimated for the latter variable directly represented the marginal utility of income (i.e., the disutility of losing money), which was used to calculate changes in economic welfare perceived by anglers at equilibrium for each lake and in an aggregated fashion for the landscape following standard economic theory (Hahnemann 1984; for an application to angling, see Dorow et al., 2010). Economic welfare relates to the notion of well-being by anglers as demand; it is a more suitable concept to economically rank policy options in recreational fisheries studies than the notion of supply that is focused on provision of fishing opportunities, such as catch rates. This is because such a supply perspectives neglects all other components of angler utility and well-being other than catch, including spatial aspects related to the location of lakes in a landscape (Cole & Ward, 1994). Put simply: a high catch rate fishery maintained close to home produces more benefits to anglers than the same catch rate offered in remote locations (Cole & Ward, 1994), and this difference in utility can only be measured by the welfare concept, not by catch rates. Note how previous landscape models have measured the catch-based fishing quality in separate “travel zones” or “regions” in the landscape (Mee et al., 2016; Parkinson et al., 2004; Post et al., 2008; Wilson et al., 2016), which conceptually controls for the disutility of travel. Still such research strictly speaking only integrates costs, catch rate and size of fish (as components of fishing quality) as generating utility to anglers. Our approach differs as the utility of a given lake is a function of multiple catch- and non-catch related utility components (harvest regulations, size of fish, catch rate, distance, cost, crowding). Most importantly the regional-level utility at equilibrium across all lakes therefore becomes an emergent property of fish-angler interactions and not one that is assumed a priori as done in related work (Parkinson et al., 2004).

Economic welfare captures the integrated nature of utilities (benefits) offered by fishing opportunities and hence represents a measure of social yield (Johnston et al., 2010, 2013, 2015). Note that economic welfare is always a relative measure of well-being emerging from a policy option A compared to some status quo or a policy option B (Cole & Ward, 1994; Fenichel et al., 2013b), i.e., welfare is assessed at the “margins”. We applied such a welfare perspective, rather than potentially incomplete surrogate such as experienced catch rates or catch-based “fishing quality”, to model runs with and without one-size-fits all harvest regulations to examine the change in regional level angler welfare stemming from regulations and the resulting changes in all lake-specific and utility-determining attributes of the experience directly or indirectly caused by regulation changes (Fig. 1, Welfare measure). The change in welfare was approximated by the change in the sum of anglers’ lake-specific willingness to pay (WTP) for a particular scenario compared to the baseline scenario and was represented in monetary units (€) (Hahnemann, 1984, equations 17 and 18 in Table 1).

We choose the no regulation scenario as the baseline and used alternative scenarios for harvest regulations to evaluate change in WTP when the common set of harvest regulations was introduced in the model. Because the marginal change in income was represented by the utility loss of annual license cost, the change in WTP (*Z_i_* of equation 18 in Table 1) represented the average change in the angling quality of angler per year for the angling quality in the entire region, i.e., welfare was a regional-level outcome metric. To relate our work also to previous catch-rate utilities, we also kept track of regional effort shifts and catch rates where needed to address our objectives.

From an ecological perspective, we estimated additional commonly used regional-level biological/ecological outcomes (Fig. 1, Conservation measures). Two outcome criteria were used to represent the status of exploited stocks at equilibrium. We chose these criteria because they were common single-species stock assessment reference points used for indicating overfished status (Worm et al., 2009). Accordingly, we defined a pike population in a given lake to be overexploited (i.e., recruitment overfished) when its spawning stock biomass (SSB) was less than 35% of its pristine, unexploited SSB (Allen et al., 2009; Mace, 1994). We considered the pike stock in a given lake collapsed if its SSB was less than 10% of its pristine SSB following Worm et al. (2009) and Hunt et al. (2011). We aggregated the number of exploited or collapsed stocks over the region, to represent regional-level conservation outcomes.

### Outline of analysis

Numerical simulations were carried out for a parameter set chosen (Tables 1, 2) to describe size-selective recreational fishing on spatially structured pike stocks by regionally mobile anglers with and without the presence of one-size-fits all harvest regulations. Similar to Hunt et al. (2011), we conducted discrete annual time-step simulations for each management scenario at a particular size of the angler population for a given residential pattern until the fish and angler populations reached a dynamic equilibrium after about 150 years. We used 10 different randomized patterns of the lake distribution and calculated an average of the 10 patterns as a value representing each simulation run.

In the simulations, we tested several scenarios or varied several variables systematically (elements shown in grey in Fig. 1). When we introduced our welfare measure, we considered two sets of harvest regulations: a no regulation case and a traditional one-size-fits all harvest regulation scenario to correspond with typical situations in many freshwater fisheries landscapes and to represent extremes. In the traditional one-size-fits all regulation scenario, we used a combination of a minimum-length limit of 50 cm and a daily bag limit of 3 pike per angler day, which is common in Germany (Arlinghaus et al., 2010) and some areas in North America (Paukert et al., 2001). In all simulations, we systematically varied the size of the angler population, which we call potential regional angling effort (to distinguish it from the realized angling effort, which is an emergent property of fish-angler interactions locally and in the region; usually only 40–60 % of the potential is realized effort). We ran simulations with and without the presence of ecological heterogeneity, with and without the presence of angler heterogeneity (by either assuming the 1-class or the 4-class angler models, Table 3) and for varying attributes of lake heterogeneity (varying the slope of the stock-recruitment function or the carrying capacity) while systematically varying the angler population size because the latter has been found before to strongly affect regional patterns of overfishing (Hunt et al., 2011).

We evaluated regional level outcomes at the dynamic equilibrium by examining both conservation objectives (SSB) as well as social and economic objectives (biomass yield, angler welfare and occasionally catch rates). Although angler welfare integrated catch rates endogenously, we singled out catch rates at equilibrium across lakes to systematically assess catch-based IFD assumptions commonly expressed in landscape studies of freshwater recreational fisheries (Mee et al., 2016; Parkinson et al., 2004).

## Results

### Objective 1 – the residential pattern shapes the geographical location of effort and overfishing, but not overall frequency of overfished stocks

When lakes were homogenous in their ecology and the (heterogeneous) angler population lived in one central urbanity in the landscape, the spatial distribution of angling effort (Fig. 2 second row) and lake-specific overfishing (Fig. 3 second row) systematically spread from the urban centre towards the periphery of the landscape as the potential regional angling effort density (AED) increased. Note that the overall level of the potential AED the landscape could support was strongly affected by the presence (Figs. 2 and 3) or absence (Figs. S2 and S3) of harvest regulations in place: harvest-regulated landscapes required much larger potential AED before the stocks collapsed entirely. In the urban landscape, the domino-like spread of overfishing from the central urbanity to the periphery was largely similar in ecologically homogenous and ecologically heterogeneous lake landscapes when lake heterogeneity was represented either by variation in productivity or variation in carrying capacity in relation to the underlying pike stock-recruitment relationship (presence of regulations, Figs. 2 and 3, absence of harvest regulations, Figs. S2 and S3). In both cases, lakes near the metropolis attracted more angling effort than more remote lakes unless regional fishing effort became excessively large for fish populations to withstand the angling pressure (Figs. S9–11).

**Figure 2.**
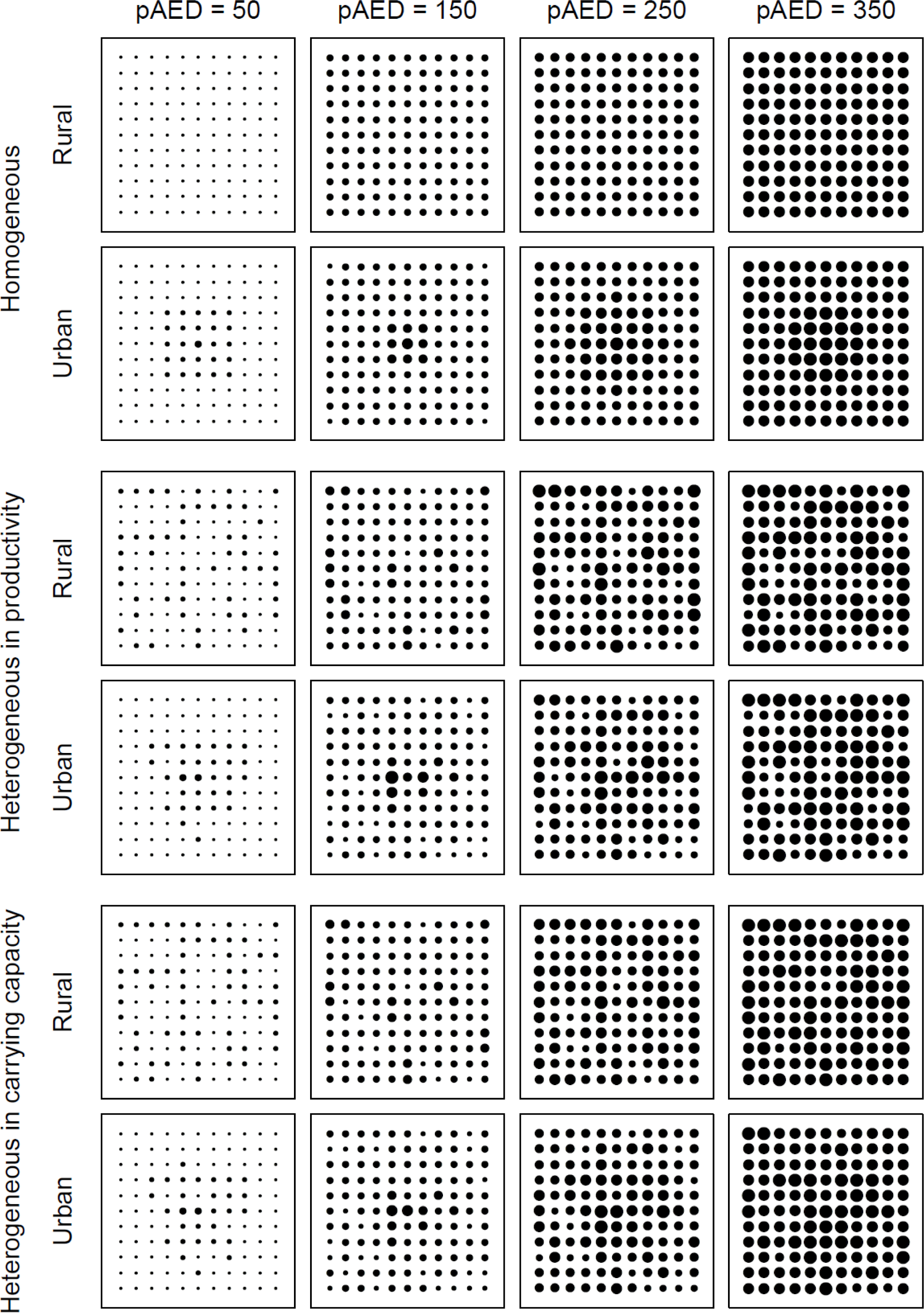
An example of the distribution of lake-specific angling effort in the homogeneous and heterogeneous landscapes with the presence of the one-size-fits all regulation in the rural and urban landscapes. Lakes are identical in the homogeneous landscape, while lakes differ in their productivity or carrying capacity in the heterogeneous landscape. The annual angling effort densities are: <30, <60, <90, <120, <150, and ≥150 [h ha-^1^].

**Figure 3.**
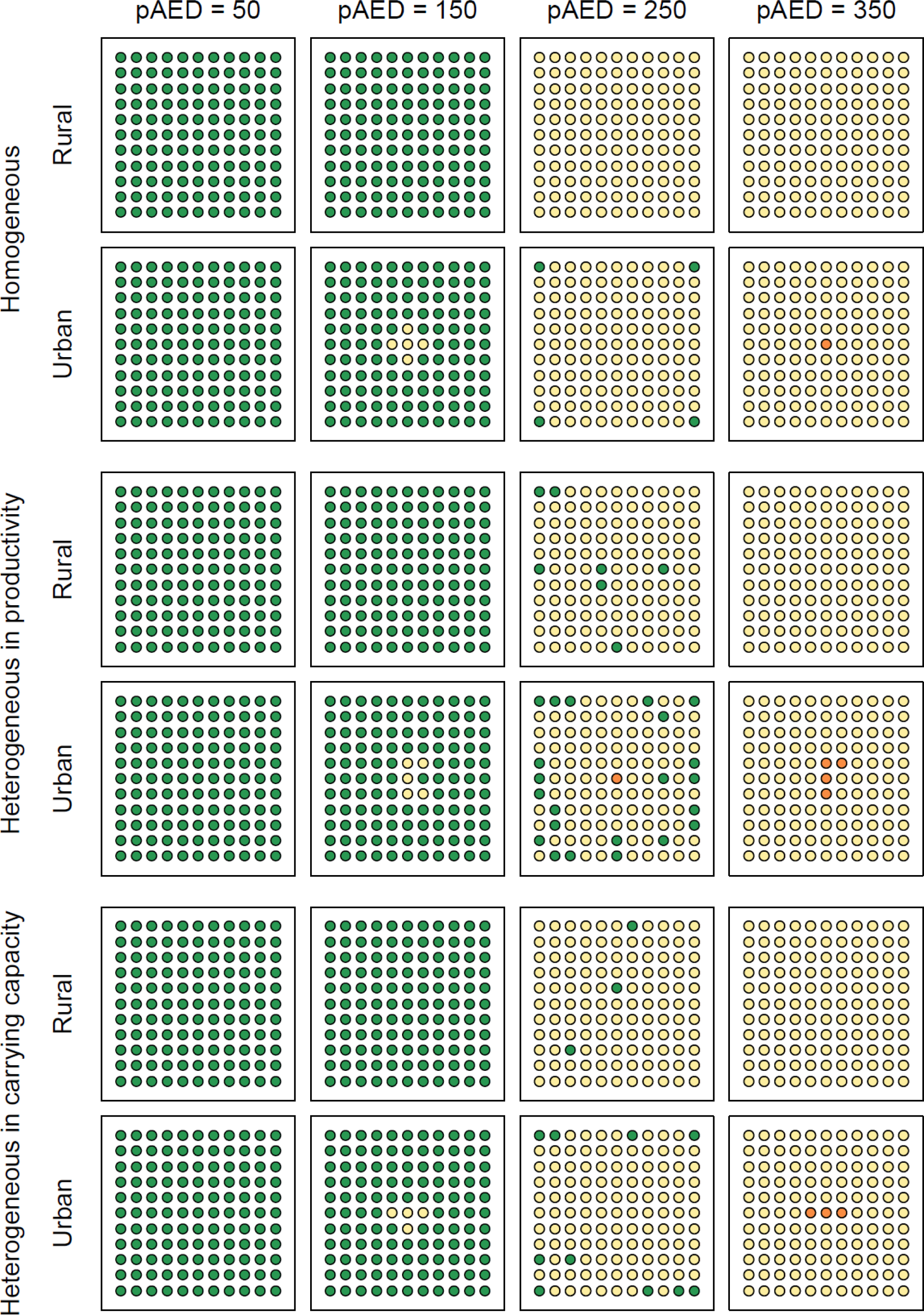
An example of the spatial pattern of exploitation in the homogeneous and heterogeneous landscapes with the presence of the one-size-fits all regulation in the rural and urban landscapes. Lakes are identical in the homogeneous landscape, while lakes differ in their productivity or carrying capacity in the heterogeneous landscape. Lakes are categorized based on their relative spawning stock biomass (SSB) to their pristine SSB (SSB/SSB_0_). Green: healthy (0.35 or higher), yellow: overfished (between 0.35 and 0.10), red: collapsed (less than 0.10). pAED is potential annual angling effort density [h ha^-1^].

The spatial pattern of lake-specific angling effort densities and overfishing at equilibrium was different in the rural landscape (Figs. 2 and 3 first row) compared to the urban landscape (Figs. 2 and 3 second row), particularly in relation to the distribution of angling effort (Fig. 2). Compared to the urban case, in the rural landscape scenario there was a much more uniform geographic placement of angling effort (Fig. 2, Fig. S2) and overfishing (Fig. 3, Fig. S3). In the rural landscape the lake heterogeneity in productivity and in carrying capacity also exerted more influence on effort density patterns and regional-level overfishing than in the urban case when comparing outcomes to the homogenous lake ecology. These effects of the rural spatial structure were particularly pronounced in the one-size-fits all policy scenario (Figs. 2 and 3) compared to the no-regulation case (Figs. S2 and S3). In general, lakes with greater potential for generating high catch-rate fisheries systematically attracted more effort, but the effect was much stronger in relation to variation in the slope of the stock-recruitment curve (productivity) than in variation of the carrying capacity (see Objective 3 below for details).

The analysis so far suggests that the location of attracted effort and overfishing is strongly driven by the potential AED (representing the size of the regional angler population in relation to available fisheries) and the residential pattern. By contrast, the aggregated regional-level outcomes of fish stock-angler interactions in terms of number or the fraction of overfished stocks, the average regional biomass yield (kg of pike per ha per year), and in the case of comparing a regulated landscape to an unregulated case also angler welfare gains, were found to be largely independent of the residential pattern or the presence or absence of lake heterogeneity both in the one-size-fits all harvest regulation (Fig. 4) as well as in the no-regulation scenario (Fig. S4). It was also largely irrelevant for overall landscape patterns of overfishing, which particular feature of lake heterogeneity varied in space (productivity vs. carrying capacity, Fig. 4). What overwhelmingly drove overall landscape outcomes was merely the size of the regional angler population in relation to available fishing area, i.e., potential AED, which often led to a realized effort to be less than 50% of the potential AED (Fig. 4, Fig. S4). For the parameter set we choose, in the no regulation case, a potential AED of about 80-90 angling-h ha^-1^ led to regional-level maximum sustainable yield (MSY), but also to a sizable fraction of about 20–40% of recruitment-overfished stocks under regional-level MSY (Fig. S4). Note that the fraction of overfished stocked rapidly increased when the potential AED moved from 80 to about 110 angling-h ha^-1^, and correspondingly the regional-level yield dropped, suggesting that a management strategy focused on regional-level MSY may render the system vulnerable to overfishing. There were corresponding trends in the regulated landscape, albeit at higher potential AED levels because the populations were better protected from overharvest (Fig. 4). Relative to the no-regulation case and at identical potential AED, one-size fits all harvest regulations led to more realized effort attracted to the landscape, a reduction in the number of overexploited lakes and maintenance of higher regional yield, which also held at large potential AED values (Fig. 5). In contrast to the biomass yield, average angler welfare constantly rose with increasing potential AED in the regulated landscape (Figs. 4 and 5). This finding was caused by the poor state of fishing in the unregulated case in the absence of regulations (Fig. S4) used as a baseline to estimate welfare gains (Fig. 4). Therefore, as a regional-level metric, angler welfare does not show a maximum that may be used as a management target (Figs. 4 and 5) as long as the unregulated case is used as a baseline. By contrast, regional MSY followed dome-shaped patterns typical for exploited fish populations in single lakes and thus maybe used as a regional management objective among others.

**Figure 4.**
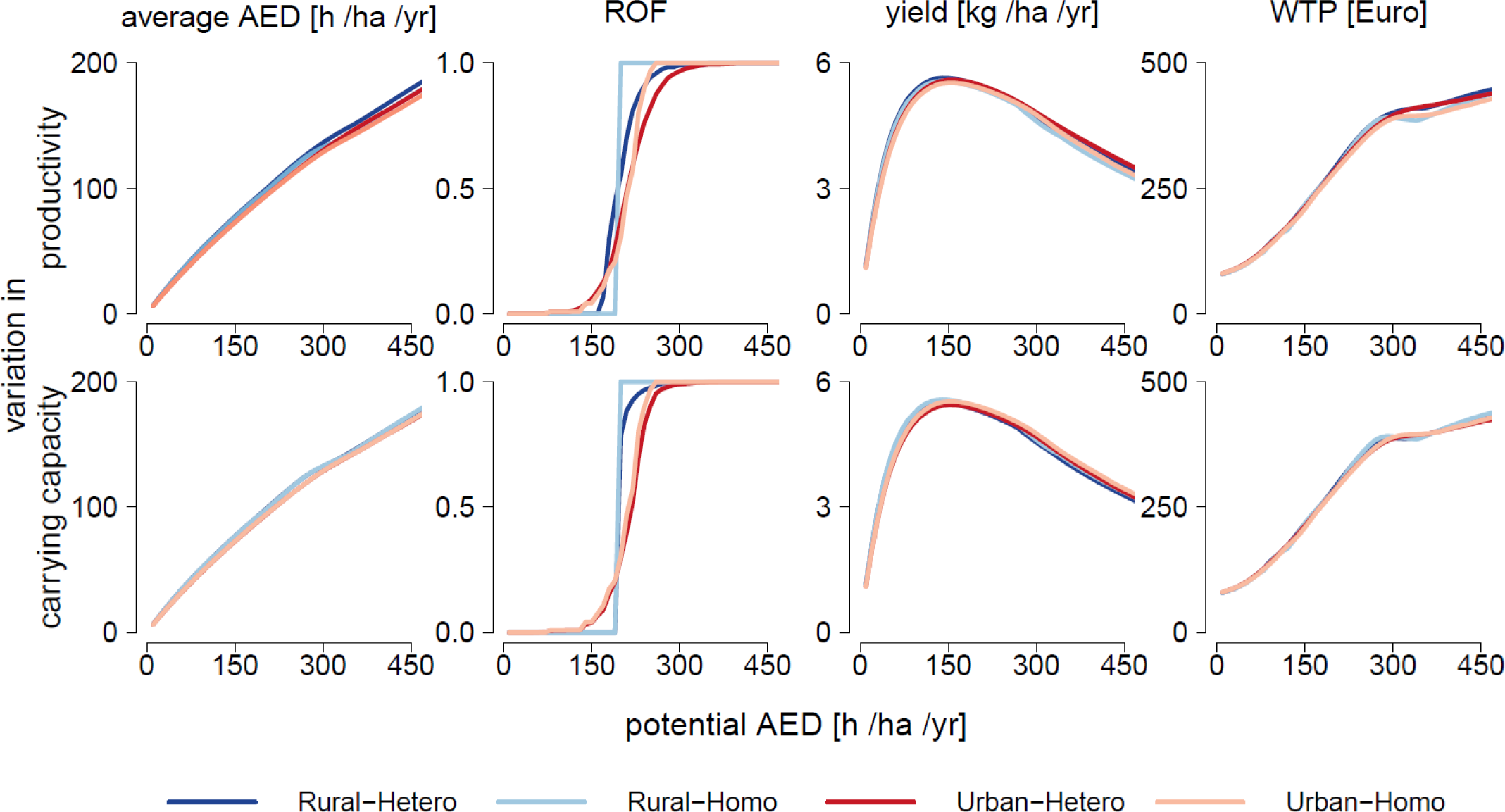
Comparison between the homogeneous (Homo) and heterogeneous (Hetero)landscapes. Lakes are identical in the homogeneous landscape, while lakes vary in theirproductivity (top) or carrying capacity (bottom) in the heterogeneous landscape. Regional outcomes in terms of average lake-specific angling effort, degree of overexploitation of lakes (ROF = recruitment overfished stocks), biomass yield, and angler welfare as represented by average willingness-to-pay (WTP) per year in the rural and urban landscapes with the presence of the one-size-fits all harvest regulation are shown.

**Figure 5.**
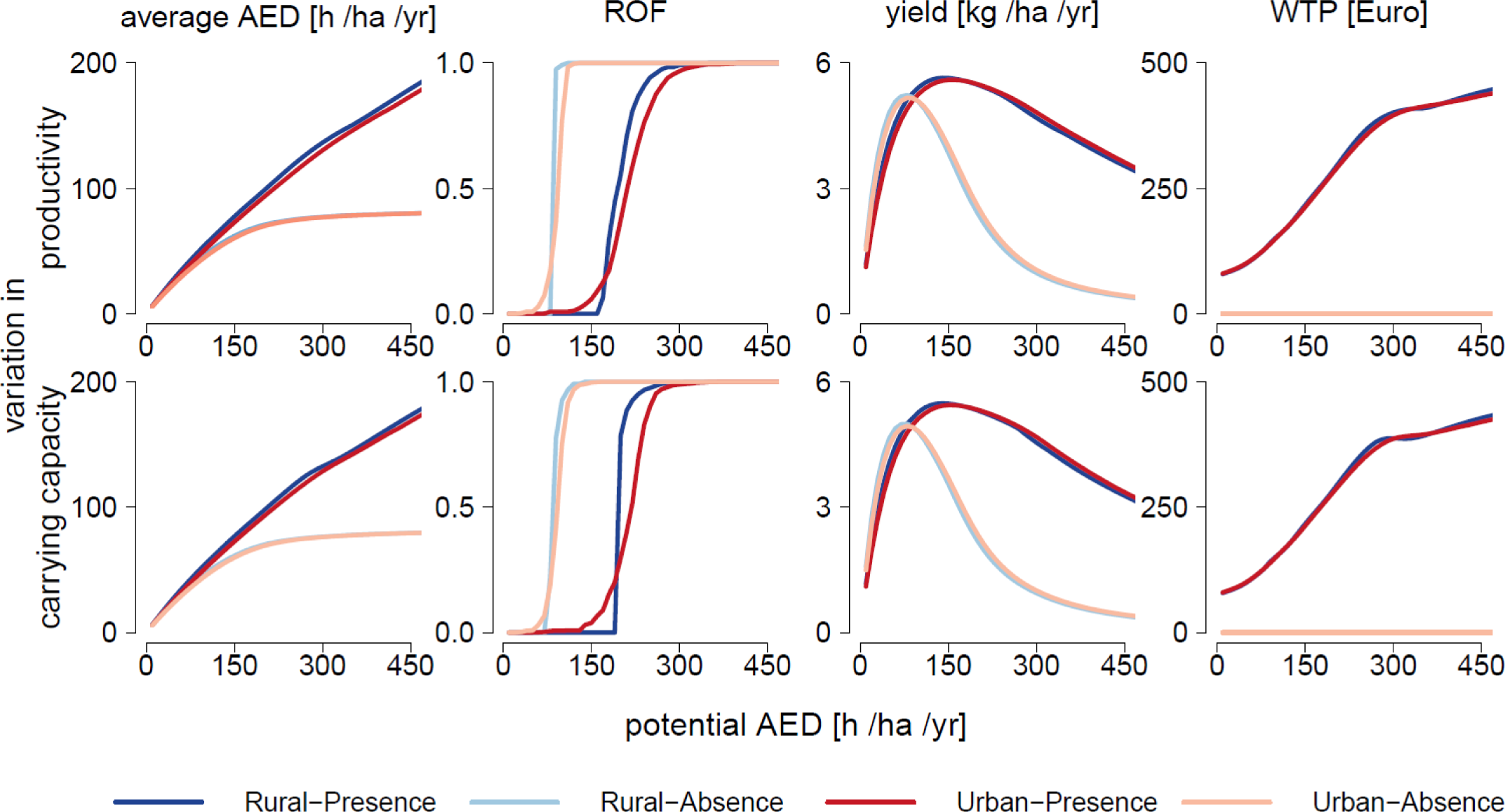
Comparison between the situations with or without the presence of the one-size-fits all harvest regulation in the rural and urban landscapes. Regional outcomes in terms of angling effort, overexploitation of lakes (ROF = recruitment overfished stocks), biomass yield, and angler welfare as represented by average willingness-to-pay (WTP) per year are shown. Lakes vary in their productivity (top) or carrying capacity (bottom).

### Objectives 2 – heterogeneous anglers exert greater cumulative fishing pressure in the region than homogenous populations of anglers

When we assumed an average empirically grounded angler type estimated from the same choice data for German anglers, we found quite different ecological and social outcomes compared to when we assumed heterogeneous anglers in the model. Figure 6 presents the results for a one-size-fits-all harvest regulation policy, and the corresponding unregulated outcomes of angler heterogeneity are shown in Figure S5. The number of overexploited lakes predicted in the 1-class model (homogeneous angler model) was always smaller than the number of overexploited lakes predicted in the 4-class model (heterogeneous model). One important contributor was the difference in the realized AED, which was always higher when multiple angler types exploited the regional fishery (Fig. 6). The maximum average regional yield did not differ between the 1-class and 4-class models (Fig. 6) because MSY was caused by purely biological properties of the fish stock. However, as the angler population size increased the total regional yield was predictably smaller in the 4-class model because the diverse anglers exerted greater harvesting pressure (i.e., realised angling effort) at the same potential AED compared to homogenous anglers.

**Figure 6.**
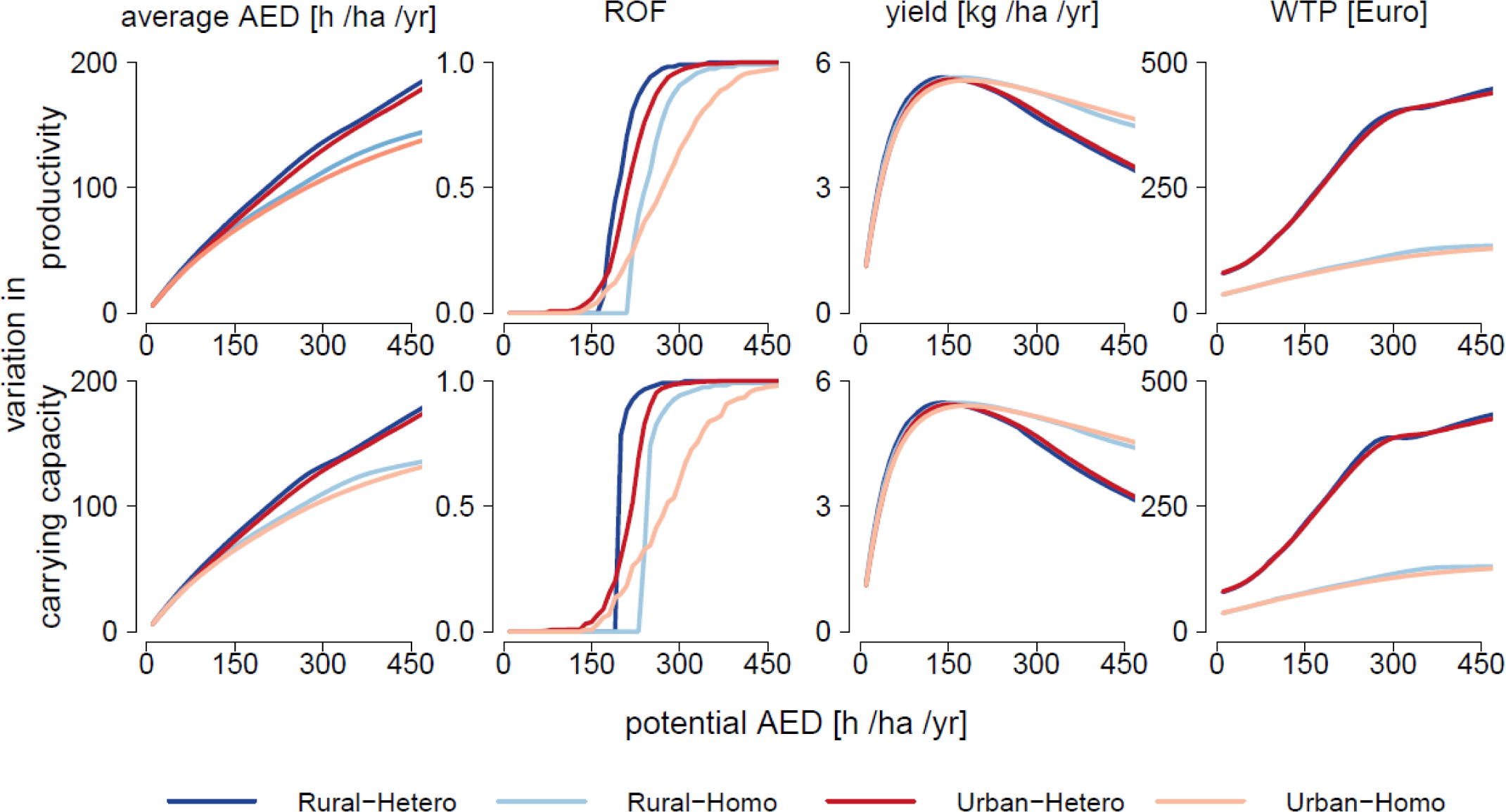
Comparison between the 4-class heterogeneous (Hetero) and 1-class homogeneous (Homo) angler models in the rural and urban landscapes. Regional outcomes in angling effort, overexploitation of lakes (ROF = recruitment overfished lakes), biomass yield and angler welfare as represented by average willingness-to-pay (WTP) per year with the presence of the on-size-fits all harvest regulation are shown. Lakes vary in their productivity (top) or carrying capacity (bottom).

The aggregated regional welfare of anglers as measured by WTP change from the unregulated to the regulated landscape was substantially greater in the 4-class model compared to the 1-class model. One large contributor to this effect was the more depressed baseline overfishing state at high potential AED in the unregulated landscape (Fig. S5) because the degree of overfishing caused by heterogeneous anglers was much more severe compared to the state of overfishing caused by homogenous anglers. Correspondingly, the welfare gains of regulations were appreciably higher for heterogeneous anglers compared to homogenous anglers. The difference in ecological and social regional outcomes among homogenous and heterogeneous anglers increased as the angler population size increased, but there was very little impact of residential patterns on regional-level outcomes stemming from the presence or absence of angler heterogeneity (Fig. 6).

The above mentioned effects of angler heterogeneity were caused by a complex pattern of spatial lake substitution patterns as a function angler preferences interacting with ecological processes of fish stock renewal. Because residential patterns did not matter much for determining the overall regional-level effects of angler heterogeneity (Fig. 6), we confine our example of where specific angler types were fishing in the landscape in the regulated urban case where we separate different travel zones of interest from the metropolis (Fig. 7, see Fig. S6 for the unregulated case). In line with our empirical data from northern Germany, the angler class 1 (committed anglers, supplemental material) made up 51.4% of the entire angler population, but this class accounted for a disproportionally larger proportion of the total angling trips taken by the angler population as a whole. The proportion of class 1 anglers in the total angling effort increased as the distance from the metropolis increased (Fig. 7) because class 1 anglers enjoyed less disutility from travel distance. By contrast, the angler classes 2 and 3 (active and casual anglers) preferred angling in lakes nearby their residence and thus rarely visited remote lakes (in zones 3 and 4 in Fig. 7). When an average type of angler was assumed instead (right panels in Fig. 7), the realised angling effort was overall lower than in the heterogeneous angler model (left panels in Fig. 7). This is because the average angler did not visit the remote lakes in travel zones 3 and 4 as often compared to the numerically dominant class 1 anglers in the heterogeneous model. The difference became more pronounced when the angler population size (potential AED) increased and the corresponding angling quality decreased because this elevated the visits to remote lakes by highly committed class-1 anglers in the heterogeneous population (Fig. 7). We can conclude that regional variation in the residency of different type of anglers will exert complex effects on landscape-scale social and ecological outcomes.

**Figure 7.**
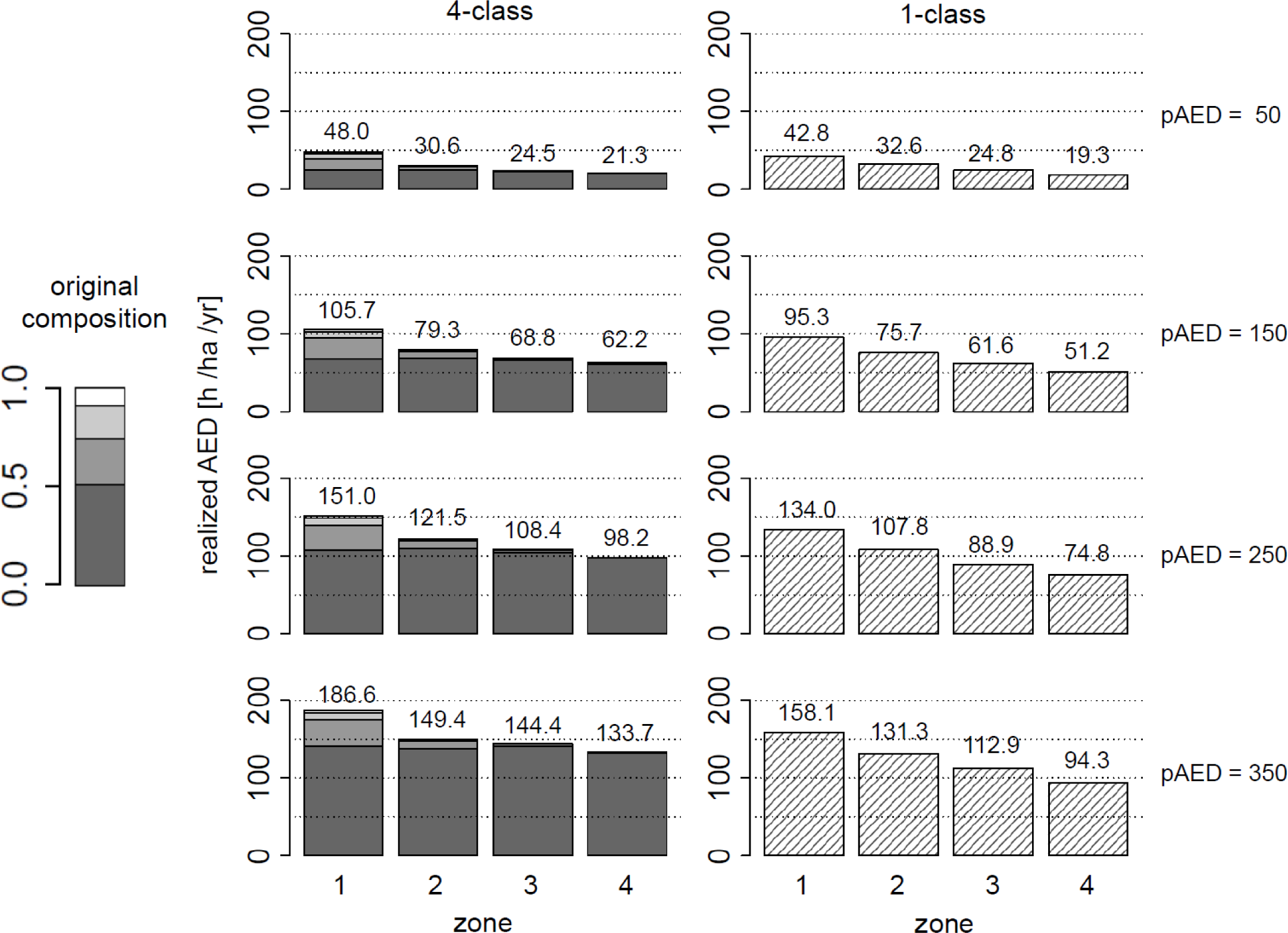
Proportions of each angler class within the realised angling effort density (AED, angling-h ha^-1^) in the urban case with the presence of the one-size-fits all harvest regulation. Lakes vary in their productivity. Lakes are categorized by the distance from the metropolis: Zone 1 (<28 km), 2 (<56 km) 3 (≥84 km) and 4 (84 km). The original proportion of theangler classes is shown on the left.

### Objectives 3 – ecological variation in production maintains catch variation unless the angler population size is excessive and lakes vary in carrying capacity not productivity

When lakes differed in their carrying capacity in the absence of regulations and were exploited by a large heterogeneous angler population, lakes of higher intrinsic quality (meaning lakes that could maximally host more fishes) tended to be exploited more heavily than lower-quality lakes, as can be inferred from a larger drop in SSB/pristine SSB as pristine SSB levels increased in both rural (Fig. S7b) and urban landscapes (Fig. S8b). In other words, positive correlations between the lake quality and degree of exploitation were found, in particular, when the angler population size was large in all landscape types (Figs. S7b and S8b). As the regional angler population size increased, the difference in the catch rates offered by the lakes in the landscape at equilibrium decreased leading to regional-level homogenization of catch rates among lakes across all lakes varying in carrying capacity in both rural (Fig. S7b) and urban landscapes (Fig. S8b).

The landscape pattern of exploitation at equilibrium differed when lakes varied in their productivity at low pike population size (slope of the stock-recruitment curve) instead of the carrying capacity. Compared to lakes varying in carrying-capacity (Figs. S7b and S8b), more productive lakes were exploited less heavily than low-productive lakes, and a homogenization of the exploited SSBs relative to pristine SBB across the productivity gradient, rather than a homogenization of catch rates, emerged as the potential AED increased in both rural (Fig. S7a) and urban landscapes (Fig. S8a). This is in contrast to the inverse relationship among pristine SSB and the exploited SSB/pristine SSB seen before for the variation in carrying-capacity among lakes (Figs. S7b and S8b). Lake heterogeneity in productivity at low population sizes also led to the maintenance of larger catch rates in highly-productive lakes at equilibrium compared to low productive lakes in rural (Fig. S7a) and urban landscapes (Fig. S8a), which contrasted with the more consistent homogenization in catch rate across lakes in all landscape types for lakes varying in carrying capacity (Figs. S7b and S8b). Substantially more variability among lakes varying in productivity persisted in the urban case also at high potential angler densities (Fig. S8a). One reason was the systematic impact of distance on lake attractiveness (utility) to anglers, which maintained fish populations at higher levels as the distance from the metropolis increased (see urban case with no regulations in Fig. S10 compared to rural case with no regulations in Fig. S9). Overall, substantial among-lake variation at the same distance in terms of annual trips that were attracted and the catch rates offered to anglers were maintained at equilibrium when lakes differed in productivity, until the angler population became excessive leading to complete collapse (Figs. S10 and S11).

The implementation of a one-size-fits-all harvest regulations (minimum-length limit of 50 cm and daily bag limit of three pike) in all lakes in the landscape modified the association of overfishing and lake quality and the ecological and social outcomes just described (Figs. 8 and 9). However, no complete reversal of the systematic patterns of the relationships of lake heterogeneity and landscape level outcomes mentioned above was found. Instead, some of the features became more pronounced. Overall, the effect of the harvest regulations was most strongly observed in higher-quality lakes than in lower-quality lakes (Figs. 8 and 9). In particular, the difference in the expected catch rates at equilibrium among high-quality and low-quality lakes became more pronounced under harvest regulations, with more productive lakes and lakes with higher carrying capacity generally offering higher catch rates than less productive lakes or lakes with lower carrying capacity in both rural (Fig. 8) and urban landscapes (Fig. 9). The positive correlation between variation in productivity and catch rate was more pronounced than that between variation in carrying capacity and catch rate (Fig. 8 and 9). In the case where variation in lake quality was arising from variation in carrying capacity, the negative correlation of pristine SSB and the exploited SSB/pristine SSB seen in the absence of regulations (Figs. S7b and S8b) was observed only when the angler population size was very large in both the rural and urban cases (Figs. 8b and 9b). Also, lakes with lower carrying capacity were only exploited more heavily than lakes with large carrying capacity when the angler population size was small and only in a rural scenario (Fig. 8b). In the case of variation among lakes in productivity this effect was even more pronounced, turning the correlation of pristine SSB and the exploited SSB/pristine SSB systematically positive across all levels of the potential AED, with no homogenization of catch rates observed among lakes (Figs. 8a and 9a, see also Figs. S11 and S12 for changes of catch rates with distance). The catch-rate homogenization was much less pronounced or not pronounced at all in the case of variation among lakes in carrying capacity when regulations were present (Figs. 8b and 9b, see also Figs. S11 and S12) compared to the no regulation case (Figs. S7b and S8b).

**Figure 8.**
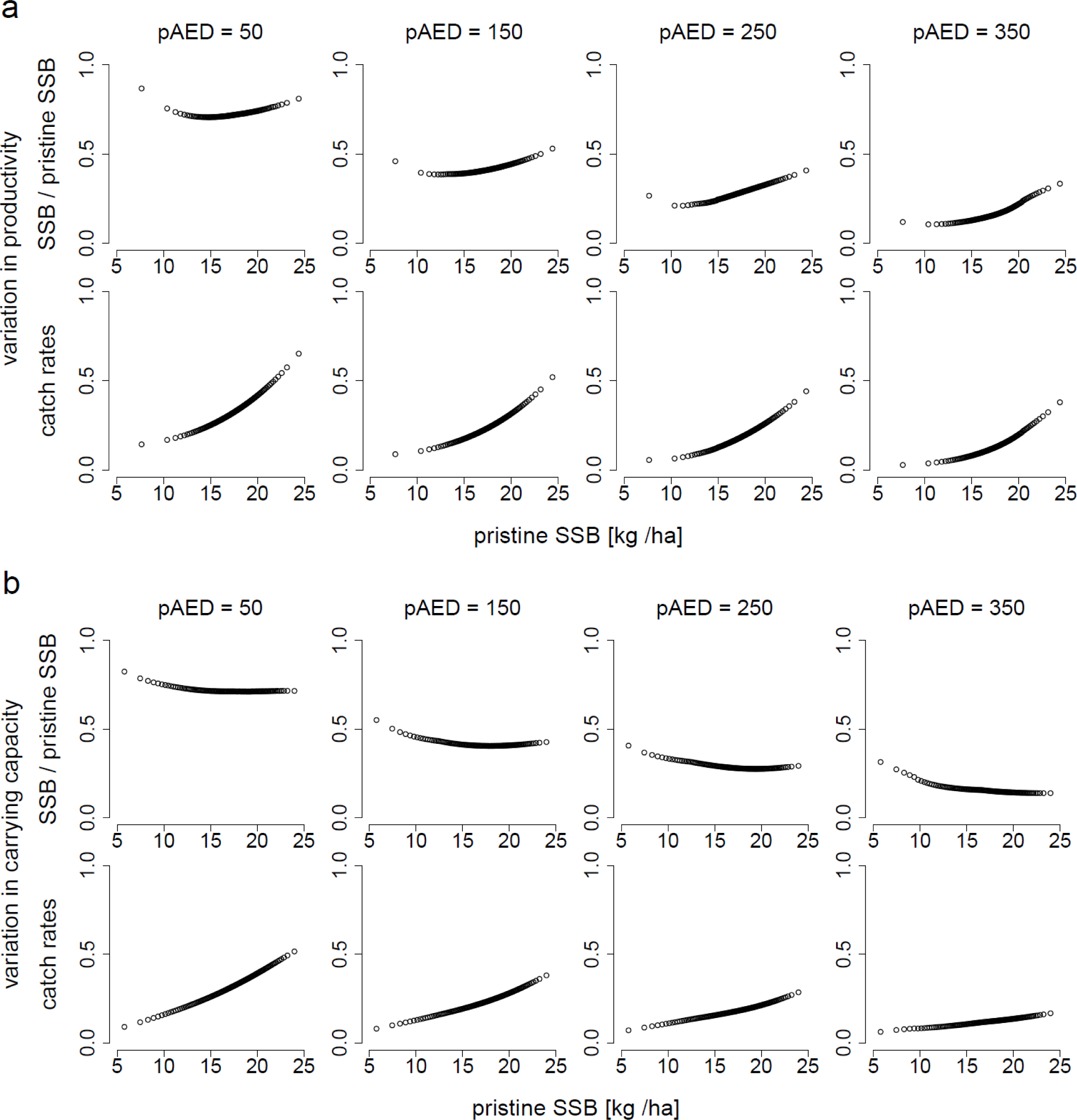
Relationship between a lake’s intrinsic quality (pristine SSB = SSB_0_) and the degree of exploitation (represented by SSB/SSB_0_) and average angler catch rates (pike per hour) at equilibrium with the presence of the one-size-fits all harvest regulation in a rural landscape. Each lake is represented by a circle. Variation among lakes in their pristine SSB arises either from variation in their productivity (a) or carrying capacity (b). From the left to the right: potential pAED (annual angling effort density) = 50, 150, 250, and 350 [h ha^-1^].

**Figure 9.**
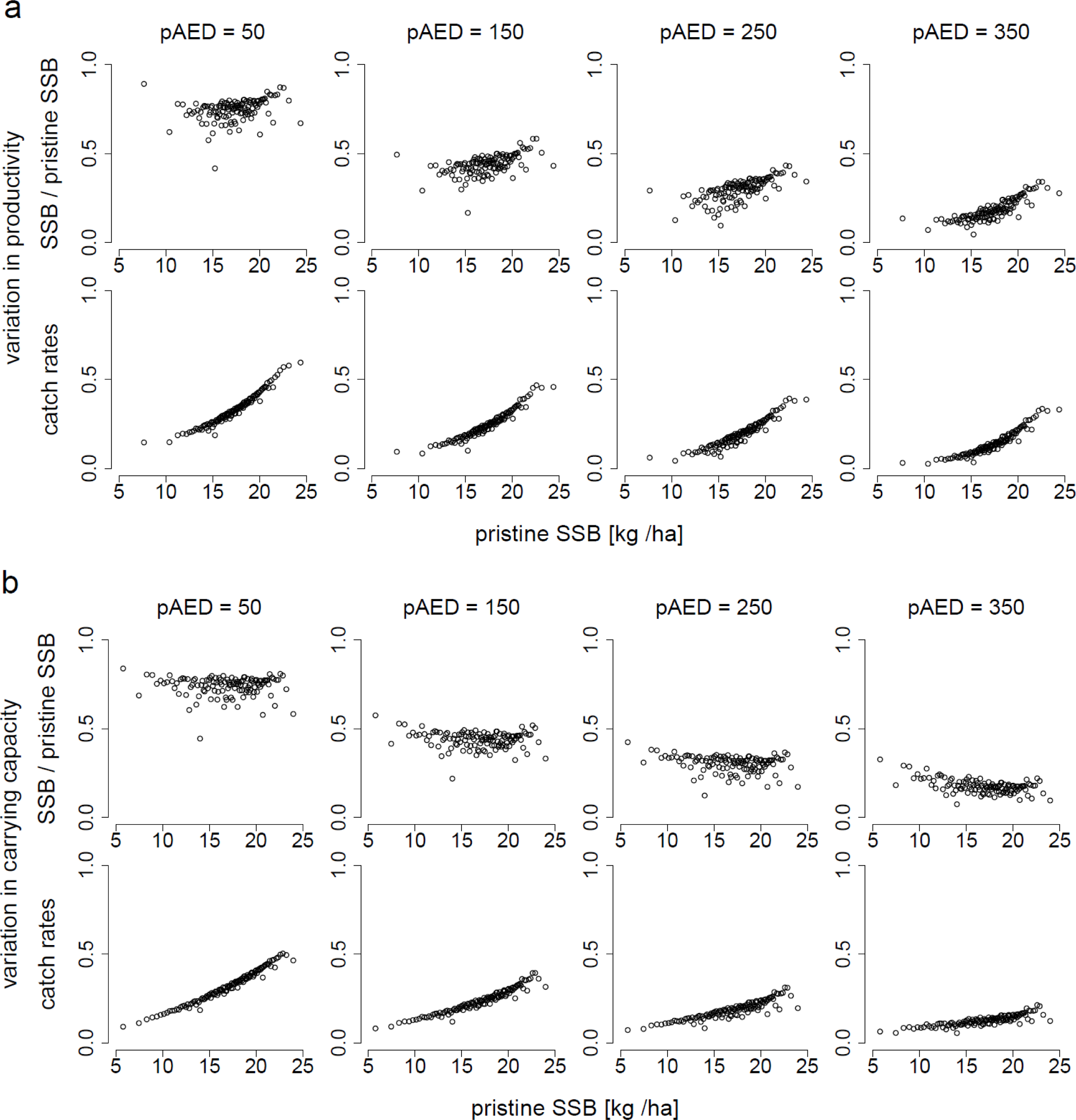
Relationship between a lake’s intrinsic quality (pristine SSB = SSB_0_) and the degree of exploitation (represented by SSB/SSB_0_) and average angler catch rates (pike per hour) at equilibrium with the presence of the one-size-fits all harvest regulation in an urban landscape. Each lake is represented by a circle. Variation among lakes in their pristine SSB arises either from variation in their productivity (a) or carrying capacity (b). From the left to the right: potential pAED (annual angling effort density) = 50, 150, 250, and 350 [h ha^-1^].

Similar patterns were observed in the urban (Fig. 9) and rural regulated landscapes (Fig. 8) in regulated landscapes compared to the no-regulation case (Figs. S8 and S7). Again, in the harvest-regulated landscape along the gradient of lake quality substantially greater among lake variability pike population size and catch rates and effort attracted was maintained in equilibrium in the rural case (Fig. 8) compared to the urban case (Fig. 9). Substantial variation in effort attracted and catch rates were present at equilibrium for lakes varying in distance in both the rural (Fig. S11) and urban regulated landscapes (Fig. S12). Variation in lakes in productivity led to somewhat greater distance-related variation in effort attracted and catch rates in both landscapes compared to variation in carrying capacity (Figs. S11 and S12).

## Discussion

We provide a general framework to examine spatial problems related to fish-stock-angler interactions and thereby contribute to an emerging literature of modelling complex adaptive social-ecological systems (Arlinghaus et al., 2017; Schlüter et al., 2012) where macro scale outcomes (e.g., regional effort distribution and overfishing) emerge from a high number of micro-level interactions (e.g., angler-fish stock interactions) (Levin et al., 2013). Our work presents the most general model for recreational fisheries published so far. It is distinguished from previous landscape models in recreational fisheries (in particular Hunt et al., 2011; Post & Parkinson, 2012; Post et al., 2008) by three key features.

First, the fish population model is age-structured, rather than being a biomass model, thereby allowing size-dependent catch expectations and the effects of size-based harvest limits to be represented; both strongly affect utility and thus effort dynamics of anglers (Arlinghaus et al., 2014; Dorow et al., 2010; Hunt 2005) and hence should be included in any realistic model of recreational fisheries (Askey et al., 2013).

Second, we used a mechanistic model of angler behaviour, predicted from an empirically estimated multi-dimensional utility function (Beardmore et al., 2013). This allowed regional outcomes to be emergent properties of model runs and avoided to investigate equilibrium conditions “forced” on the model by strong assumptions, such as the one that at an IFD equilibrium all fish stocks should be fished down to an average catch rate (Parkinson et al., 2004). Although intuitively appealing, we think there are limitations in the analogy of anglers and fish forming natural predator-prey systems because the fitness of the human predator (forager) far extends beyond resource intake rates (i.e., catch rates or other measures of catch quality) and thus its behaviour is more complex than the one of natural foragers. Previous modelling work has already shown that assuming anglers to be mainly or exclusively driven in their behaviour by catch expectations can lead to unrealistic predictions about how best to serve their expectations from a management perspective (Johnston et al., 2010). We therefore suggest our model is mechanistically superior to models that assume that human foragers are simply guided by catch expectations, unless one can show that a particular angler population is indeed mainly driven by catch (Hunt et al., 2011). Even in the recent work by Wilson et al. (2016) and Mee et al. (2016) where the trade-offs of expected numbers and size of fish were elegantly expressed using region-dependent catch quality “isopleths”, substantial among lake variation in catch qualities remained at equilibrium, suggesting more factors than catch aspects affected lake utility and in turn angler participation and effort allocation. Therefore, we suggest the null model for representing angler behaviour is one that assumes a multi-dimensional utility function composed of both catch- and non-catch attributes, rather than a fitness function exclusively driven by catch expectations.

Third, rather than focusing on just selected regional outcomes (e.g., number of overexploited stocks, Hunt et al., 2011; or fishing quality, Mee et al., 2016; Post et al., 2008), we calculated and presented several emerging outcomes jointly, which encompassed regional-level ecological (e.g., regional overfishing) and socio-economic objectives (e.g., regional angler welfare) as well as more traditional objectives of recreational fisheries (e.g., catch rates and effort). Thereby, our model accommodated important trade-offs in management objectives and associated criteria explicitly.

Our key result is that landscape patterns of overexploitation are an integrated function of angler and lake heterogeneity as moderated by residential pattern, angler population size, the type of lake quality variation (productivity or carrying capacity) and the presence or absence of harvest regulations. In terms of largely robust predictions we 1) confirmed earlier studies that in urban landscapes lakes around the aggregation of effort will receive greater effort and overfishing risk than more remote lakes (*sensu* Post et al., 2002; Carpenter & Brock, 2004; Post et al., 2008), 2) found that angler population size and angler heterogeneity aggravates the degree of overfishing by spreading effort more across lakes (similar to Johnston et al., 2010 in a single lake case and Hunt et al., 2011 in a regional case), and 3) reported that the previously proposed hypothesis that higher (ecological) quality lakes will be systematically overfished by regionally mobile anglers (Parkinson et al., 2004) and that at equilibrium all lakes (within zones of similar travel distance) will be offering similar catch rates (Parkinson et al., 2004) or catch qualities (Mee et al., 2016; Wilson et al., 2016) are confined to particular cases or empirical systems and cannot be easily generalized. In fact the positive association of lake quality and degree of overexploitation (as judged by SSB relative to pristine SSB) was only found for unregulated (be it rural or urban) landscapes at high potential angling effort when lakes varied in carrying capacity, but not in productivity. A further clear-cut result we found was that an increasing angler population size will have systematic overfishing effects and reduce both equilibrium stock sizes and average catch rates irrespective of residential pattern, lake heterogeneity and the presence of angler diversity, but unless we have extreme situations (e.g., exceedingly high potential angling effort), substantial among lake variation in expected catch rates still remained. We discuss our detailed results first with reference to the three objectives stated in the introduction before moving to model limitations and implications for management and policy making.

### Discussion of the three principal objectives

The first key finding of our modelling experiment was that the spatial patterns of angling effort attracted and regional overfishing were dependent on the residential patterns in a given landscape as moderated by the angler population size and was less affected by ecological heterogeneity among lakes. Our work agreed with previous landscape models reporting that overfishing of spatially structured fish stocks proceeds in a systematic fashion from aggregation of high latent angler effort in urban landscapes towards the periphery (Carpenter & Brock, 2004; Hunt et al., 2011; Post et al., 2008), and we found this pattern was not strongly affected by lake heterogeneity in urban environments. At equilibrium urban environments also maintained greater among-lake variation in expected catch rates compared to rural case because urban environments always offered some effort “refuges” in lakes in remote localities. Such effects were not present in rural landscapes, and even in an urban landscape domino-like overharvesting at high angler population sizes did not occur when the landscape was regulated by harvest regulations, supporting earlier work by Hunt et al. (2011) and Post and Parkinson, (2012).

Results from urban landscapes have so far dominated the literature on freshwater fisheries landscapes (e.g., Hunt et al., 2011; Post et al., 2008). We show that findings from urban cases do not hold for rural landscapes in relation to the spatial arrangement of overfished stocks when the regional angler population is moderate or low. That said, aggregative metrics of regional-level outcomes, e.g., the total number of overfished stocks, were found to not strongly deviate in urban and rural landscapes and be less affected by lake heterogeneity, suggesting that when the aim is to outline broad-scale outcomes simulation of urban landscapes may prove suitable approximations independent of exact knowledge of local-level productivity of ecosystems.

In relation to our second objective we can conclude that simplifying a heterogeneous angler population to a homogenous one, or to aggregates such as “angling effort”, in modelling experiments risks severely underestimating landscape-level realized effort and regional overfishing and also strongly affects the location to which effort (and overfishing risk) is attracted. This finding agrees with recent literature reviews who noted that being explicit about which behavioural responses to expect is crucially important for understanding and managing recreational fisheries (Arlinghaus et al., 2017; Ward et al., 2016). Moreover, not accounting for angler heterogeneity in preferences in behaviour underestimates the social welfare gains from harvest regulations and thus also bears strong relations to economic and managerial dimensions (Cole & Ward, 1994). Our work confirms single-lake bio-economic models in recreational fisheries showing that accounting for variation in angler types through the integrated nature of multi-attribute angler utility is important for inferring fish population developments and identification of optimal input and output regulations that maximize benefits to anglers and minimize ecological impacts (Johnston et al., 2010, 2013, 2015). Hence, it is not only of narrative importance of being explicit about which angler typologies, and relatedly variation in preferences and behaviour, exist in a given SES of recreational fisheries if the aim of the modelling experiment is to provide robust insights for management (*sensu* Cole & Ward, 1994; Fenichel and Abbott 2014; Johnston et al., 2010; Post et al., 2008). Our finding about the large importance of angler diversity for outcomes constitutes a relevant innovation because all previous landscape models of recreational fisheries have either assumed various scenarios of homogenous anglers (that vary by importance attached to catch vs- non-catch utility components, Hunt et al., 2011) or have aggregated effort of all angler types jointly (Camp et al., 2015; Post et al., 2008), sometimes further separate by “travel zones” that control for the systematic effort sorting effect caused by angler variation in accepting travel costs for the benefits of accessing lakes offering high utility (Mee et al., 2016). We think that future studies are well advised to be more explicit about which angler type the model is designed to represent, and we suggest that the angler specialization framework is particularly suited to address angler heterogeneity (Bryan 1977; Johnston et al., 2010). Different angler types not only differ in their travel propensity, but may also strongly differ in their skill and catchability (Johnston et al., 2010; Ward et al., 2013a,b), which we did not explicitly model. Further work on the relationship of angler preferences and skill/catchability is needed to improve the modelling of angler heterogeneity on landscapes.

In relation to our third and last objective, we confirmed previous studies (in particular Hunt et al., 2011) that the assumed positive correlation among exploitation impact and the ecological quality (productivity and carrying capacity) of a given lake (Parkinson et al., 2004) is to be expected only under very particular conditions and is by no means a general result. By the same token, according to our work and others (Hunt et al., 2011; Matsumura et al., 2010), a catch-based IFD where the lake-level catch rates, or more generally catch-based fishing qualities (Mee et al., 2016; Wilson et al., 2016), are homogenized across a region is not to be generally expected in recreational fisheries. In fact, based on our model we claim that the systematic overexploitation of high quality fisheries should not be expected as a default, and we also found limited evidence in our model for systematic homogenization of catch rates across lakes. These results agreed with previous modelling studies that also used a multi-dimensional utility function driving angler behaviour (Hunt et al., 2011) or assumed suboptimal patch choices of foragers (Matsumura et al., 2010) similar to the way we represented site choice behaviour of human foragers in our model. We are thus confident that our findings about departures of catch-based IFD relate to the mechanistic assumption that the fitness of the human forager relates to multiple dimensions, both catch- and non-catch related, and that human foragers suboptimally and probabilistically choose lakes offering the highest utility. Landmark work by Wilson et al. (2016) and Mee et al. (2016) that report that anglers homogenize catch qualities to size-number quality isopleths in regions differing by travel costs from the urban environment of Vancouver in fact “control” for three key dimensions of utility to anglers (size, catch rate and distance). Although some form of IFD was found in their stocking-based rainbow trout fisheries, substantial among lake variation in equal travel zones remained, most likely because other aspects than those measured affected angler utility and hence site choice.

In our study, the strongest evidence for a systematic overexploitation of high quality lakes and for catch-rate homogenization effects across both the lake quality and distance gradients in urban cases was revealed when the variation in ecological lake quality was caused by lake heterogeneity in carrying capacities in the absence of harvest regulations and for very large (and heterogeneous) angler population sizes (Figs. S7 and S8). When harvest regulations were present, however, these effects were only present at exceedingly high angler population sizes (Figs. 8 and 9). By contrast, when lakes varied in their productivity, more productive lakes were less heavily exploited and they also maintained larger catch rates compared to low quality lakes (Figs 8, 9, S7 and S8). Our results thus appeared to contradict the idea that homogenization of catch-based fishing quality or catch rates across fisheries landscapes in zones of equal access costs (Mee et al., 2016; Parkinson et al., 2004). However, this is not the case. Based on our study, for a catch-based IFD to happen, angler utility must be mainly or exclusively about expected catches, lakes need to be open to a large pool of anglers, with easy access, variation in lake quality must be based on carrying capacity, but not in the slope of the stock-recruitment curve (productivity), and no harvest regulations offering protection to the fishes should be present. Most of these ingredients apply to the stocking-reliant rainbow trout fisheries in British Columbia, for which a catch-based IFD in recreational fisheries was reported (Mee et al., 2016; Post & Parkinson, 2012; Post et al., 2002, 2008; Wilson et al., 2016). Importantly, as mentioned before in these studies a fishing-quality based IFD has been reported in travel regions varying in travel distance from the metropolis (Mee et al., 2016), which essentially controls for the systematic impact of travel on utility and site choice behaviour. Thereby, a key non-catch dimension of angler utility, distance, is removed and the angler behaviour within a given zone is in turn affected mainly by catch expectations related to catch rates and sizes of fish that are captured. The British Columbian lake systems are open-access to a large pool of anglers residing in Vancouver, they are mainly directed at harvest-oriented anglers, the lakes have few harvest regulations and variation in catches and sizes of fish to be expected among lakes is essentially a function of the stocking density as most lake rainbow trout stocks are not self-recruiting. In such situations, stocking essentially determines the carrying capacity because there is no internal renewal process at low stock sizes similar to the effects stemming from variation in the slope of a stock-recruitment curve in a naturally reproducing stock. According to our study, all these conditions indeed foster the emergence of a catch-based IFD, in line with the results from British Columbia. However, these conditions are not generally present in other fisheries landscapes, where a large fraction of fisheries are based on naturally recruiting fishes that naturally vary in productivity (i.e., slope of the stock-recruitment relationship) among systems and where at least some form of harvest regulation is present. Under such conditions, our model does not predict a catch-based IFD to easily emerge. Instead, in most landscapes the maintenance of substantial variation among fisheries in fishing utility (“quality”), rather than its erosion, is to be expected at equilibrium.

Following our model, in unregulated landscapes variation in productivity (i.e., population renewal speed) among lakes will either lead to homogenization of overfishing, while maintaining high catch rates in more productive stocks, or help maintaining both high spawning stock biomasses and high catch rates under regulated conditions in the most productive stocks. The reasons for the strongly different patterns of the SSB and catch rates in the exploited equilibrium in relation to varying carrying capacity and population renewal (i.e., productivity) in our model are purely ecological, confirming the importance of studying both ecological and social processes in coupled SES. Variation in carrying capacity will mainly lead to variation in catch rates in the unexploited state, which cannot be sustained as angling effort responds. Consequently, due to rapid effort responses of anglers at equilibrium yield, and relatedly catch rates, produced by exploited fish stocks are rather insensitive to increases in carrying capacity, and similarly variation in catch-dependent angling quality at MSY is largely independent of underlying carrying capacities of a given lake (Parkinson et al., 2004). By contrast, yield and catch-related angling quality increase strongly with increasing productivity (slope of the stock-recruitment relationship) at MSY (Parkinson et al., 2004) leading to more resilient stocks, unless they are exploited by a large pool of anglers leading to their collapse (Post et al., 2002, 2008). Hence, variation in population renewal processes at low stock sizes (i.e., productivity) can better maintain fish stocks and catch rates under exploiting conditions by compensating for losses due to fishing, which variation in carrying capacity alone cannot achieve (Walters & Martell, 2004).

By contrast, as implied by our model management interventions that modify the population renewal capacity (e.g., due to enhancement of juvenile habitat) rather than carrying capacity per se can have sustained, systematic effects on maintaining variation in catch rates and spawning biomasses in fisheries landscapes. In fact, when lakes vary in productivity rather than carrying capacity and when a base set of harvest regulations is introduced, in our model high-quality lakes become less overexploited compared to low-quality lakes. In other words, high productivity coupled with a protection of young, immature fish through a basal set of size-limits is key for lakes to avoid being systematically overfished (Post & Parkinson, 2012).

### Limitations and extensions

As any model, our work has several limitations stemming from simplification of processes and structural uncertainty. On the biological side, our work constitutes a single-species age-structured model that omits multi-species interactions in complex food webs and represents density-dependence phenomenologically rather than being an emergent property of size-structured interactions. However, pike populations exhibit a high degree of intraspecific population regulation through cannibalism and overall show stable dynamics (Persson et al., 2004). Moreover, most of the size- and density dependence was estimated from one stock (Windermere) that shows exceptionally high quality data. Obviously our model cannot be used to derive predictions for specific empirical systems, but we think that we have captured the most essential population dynamical processes in a rigorous fashion, with substantial empirical data support.

A further limitation of our work that might limit the direct comparison to other landscape models may be inherent in the different spatial scales. For example, our landscape scale encompassed 150 km, while Post et al. (2008) modelled > 1000 km, and Hunt et al. (2011) about 300 km. The reason for the different scales in the three studies relates to the calibration of the angler model, which was always empirically grounded to local conditions. Obviously, a larger scale in our model would have substantially affected the location of effort because anglers in northern Germany for which the base angler model was calibrated are not used to travel much farther than about 200 km for a single angling trip. Hence, the distance effects might have been stronger if the choice model used to construct the travel cost coefficient would have exposed anglers in the model to much larger travel distances. Modelling a landscape that is much larger than what the empirical anglers were normally exposed to would have been to extrapolate beyond the parameter space used to train the model, and hence was not done in the present study.

What might more fundamentally affect model outcomes are connections among the ecological systems, e.g., through rivers or creeks linking lakes. Newbold and Massey (2010) showed that such spatial connectivity of the fish resources affects the estimation of utility models and may demand alternative structural models of angler site choices that captures species sorting behaviour and spatially connected population dynamics. Further work in this area is certainly warranted.

We assumed that anglers were omniscient about the utilities offered by each of the lakes in the landscape and that the past year’s experiences were instantaneously exchanged among all anglers. In reality, anglers will of course not be omniscient about all lake utilities and they might also follow different strategies in terms of lake choices than we assumed. For example, rather than being utility maximizes anglers might follow different approach to lake choice (e.g., satisficing, Wierzbicki 1982). Relatedly, place attachment, habitat, tradition and the attainment of angling experience and skill with a given lake over time may all lead to “local adaptation” and a tendency for anglers to always visit familiar sites. Such effects were not included in the present model and are certainly relevant sources of uncertainty. Anglers may also strongly vary in skill (and catchability, Ward et al., 2013a), and hence the “update speed” of which catch to expect in a given lake may systematically vary among anglers, in turn affecting outcomes. All of these aspects could have strong effects on regional distribution patterns and thus need to be accounted for in future work.

We simplified our models by assuming equal skill among anglers, no social interactions between anglers other than crowding effects emerging from the utility offered by lakes, no opportunity for learning and adaptation of preferences and no social networks. All of these assumptions are unlikely to hold in any empirical system because anglers are known to differ in skill (Dorow et al., 2010; Ward et al., 2013a), are unlikely to be omniscient (Hunt et al., 2011), are characterized by shifting expectations and preferences (Gale, 1987), and quite certainly form social groups and networks of like-minded friends and peers (Hahn, 1991; Ditton et al., 1992) through which information flow happens (Little & McDonald, 2007). Such information flow changes in quality and quantity through rapid changes in novel communication technology (e.g., social media, Martin et al., 2012). Missing links among nodes in angler networks can then block information about fishing opportunities if there is strong modularization in the network (Little & McDonald, 2007), or it may foster the exploitation of lakes through slow, but steady, information spread in small world networks. The latter effect is more likely at time scales that we measured, which is why we feel confident that the lack of consideration of among-angler networks did not fundamentally bias our long-term predictions. However, anglers in networks may derive utility from the utility experienced by fellow peers (e.g., catch of a trophy by a close friend), which can affect the policy options and create sites the networks prefers as a whole (Neilson and Wichmann 2014). Also, isolated events like as the popularization of exceptional fishing opportunities may lead to a systematic and pervasive shift in effort (Carpenter et al., 1994) – a dynamic not represented in our model. All these issues are likely related to angler heterogeneity (some anglers are more networked than others, some anglers are more receptive to media than others, Ditton et al., 1992), which we found will strongly affect the location of effort and hence landscape patterns in particular empirical systems. But despite this empirically relevant complexity that should be certainly tackled in future work, we think that the long-term strategic predictions that our simple model allows may nevertheless serve as qualitative approximations of which family of outcomes to expect under particular situations.

Finally, limitations relates to omission of specific details of the governance system. We explored open-access fisheries where anglers can choose lakes in an open landscape and a social planner installs one-size-fits all policies for the entire landscape, as it typical in North America (Lester et al., 2003). However, in West Germany and many other areas of Europe small angling clubs manage restricted water areas and anglers cannot easily switch among small angling clubs (Daedlow et al., 2011), which will lead to different dynamics in the region than modelled in our paper. Again, it will be worthwhile to analyse more constrained choices and what landscape level outcomes to expect.

### Policy implications and future directions

Based on our work we can derive some management implications of potential relevance to policy makers and managers charged with managing freshwater fisheries landscapes. We outline four due to limitations in space.

First, introduction of harvest regulations following a simple one-size-fits-all policy can decrease regional overfishing and maintain high yields, while at the same time strongly increasing angler welfare compared to the unregulated case. Our work confirms earlier landscape studies that reported that to manage open-access freshwater fisheries and avoid sequential collapses some base level of regulations or other type of management intervention is necessary (Cole & Ward, 1994; Lester et al., 2003; Post & Parkinson, 2012). However, it is very likely that a diversity of management tools rather than one-size-fits all policies as examined in our model will produce better outcomes (Carpenter & Brock, 2004; Post & Parkinson, 2012). Future work is reserved for using our model to test the design (type and regional placement) of various policy and management options to optimize specific management objectives.

Second, in line with previous work (Post & Parkinson, 2012) our work implies that to achieve high regional level fish yield and avoid localized collapses of stocks, constraints on total effort (and by the same token total fishing mortality) are necessary if the latent regional angling effort is exceedingly high relative to available fishing area under open-access situations. We found that the transition from a situation that produces regional MSY to rapid collapse is narrow in terms of what average angling effort per hectare the system can maintain, which suggests that a precautionary approach may be needed to limit the total number of anglers for a given landscape if the aim is to manage for MSY. Similarly, our work and related studies suggests that if the objective is to minimize the number of overexploited stocks at high potential angling effort constraints on fishing mortality (e.g., through implementation of restrictive harvest regulations), strategic use of stocking near high aggregations of anglers (to “absorb” mobile angling effort) or even effort controls will be necessary in at least a fraction of the otherwise fully accessible ecosystems (Cox & Walters, 2002; Post & Parkinson, 2012).

Third, our model results exposed some fundamental trade-offs that mangers may need to navigate when managing mobile anglers in freshwater landscapes interacting with local ecological processes of density and size-dependent population regulation under open-access situations. In particular, in urban environments it may not be possible to maximize regional-level objectives (e.g., regional MSY) with one-size-fits-all regulations without collapsing some of the stocks in the landscape. This finding is equivalent to insights from multi-species models in the marine environment where achieving multi-species MSY comes at the cost of collapsing some stocks (Worm et al., 2009). Fully avoiding collapse when targeting regional-level MSY may only be possible in an open-access situations where anglers will always dynamically respond to changes in local fish availability by radical landscape-level effort controls, continuous stocking or implementation of total catch-and-release policies with low hooking mortality in the absence of illegal harvest (Johnston et al., 2015; Post & Parkinson, 2012). Further simulation work is needed to address this important issue.

Finally, although we did not directly model stocking-based recreational fisheries, some of our findings can be interpreted in light of previous landscape work under stocked situations, calling into question the efficiency of one-time stock enhancement activities in non-recruiting stocks when mobile anglers interact with spatially structured resource patches. As mentioned before, Mee et al. (2016) reported that mobile anglers targeting stocked rainbow trout in British Columbia quickly fish down stock-enhanced population to some regional level “fishing quality” dictated by distance-clustered number (catch)-size trade-offs for fishing quality. Similarly, in our model where the disutility of distance and the catch preferences of anglers were endogenous for the angler movement dynamic, we found that variation in lake qualities by varying the carrying capacity among lakes did not maintain variation in catch rates when the angler population size was large. In naturally recruited species elevation of the carrying capacity can only be achieved by either improvements to habitats (which is rarely implemented in practice) or through the successful stocking of usually recruited (i.e., large) fishes leading to put-and-take type of fisheries (Arlinghaus et al., 2015; Camp et al., 2017; Rogers et al., 2010; Ziegler et al., 2017). Such type of manipulation of the general availability of fishes to capture (conceptually represented by an elevated carrying capacity) is, however, not expected to have long-term effects as dynamic angling effort quickly uses any locally available utility and moves lake-specific utilities to a regional average utility offered by all lakes in the landscape. By contrast, elevating the slope of the stock-recruitment curve, for example by habitat enhancement, has been shown in our work to maintain variation in angling qualities in the region and thus could be a superior long-term strategy, knowing that variation elevates the resiliency of exploited systems (Carpenter et al., 2015). Further models on the systematic effects of stocking vs. other management options are needed because we did not explicitly model stocking in our model.

## Conclusions

We found that social and economic outcomes to be expected as emergent properties from a pool of anglers interacting with a spatially structured lake system were strongly driven by the particular spatial configuration, angler population size in relation to available lake areas and angler and lake heterogeneity. Simplification of any of these ingredients will impair the ability to predict the geographic configuration of key outcomes of interest, such as the degree of local and regional overexploitation, the angler effort attracted to specific fisheries and the well-being of fishers generated by a freshwater landscape as a whole. At the same time we also found that if one is only interested in understanding overall regional outcomes, simplification of spatial configurations and lake heterogeneity may not be overly consequential. By contrast, simplification of angler heterogeneity will lead to large biases at best, and mismanagement and stock collapses at worst. Social-ecological landscape models are one tool to systematically examine how spatial and angler heterogeneity interact with regulations to produce regional-level outcomes. Models such as ours can be an important research tool to conduct “virtual ecologist” experiments to design optimal sampling strategies and test management strategies in the framework of uncertainties using a management strategy evaluation framework (e.g., Deroba & Bence, 2008; Thébaud et al., 2014; Wilberg et al., 2008). Future work is needed engaging in multi-criteria optimization (by accounting for multiple objectives both conservation and angler well-being oriented) and how to put a landscape perspective into operation in light of severe limitations in monitoring abilities in data-poor situations (Fayram et al., 2009; Lester et al., 2014).

## Acknowledgments

Work on this project received funding with the Adaptfish program funded by the Pact of Innovation and Research by the Leibniz-Community. We thank many colleagues and team members for constructive feedback over the years. We have no conflict of interest to declare.

## Appendix I

### Estimating a generic utility model for anglers

Most previous revealed or stated preferences models of anglers were directed at a particular target species (e.g., Dorow et al., 2010; Oh & Ditton, 2006,). Beardmore et al. (2013) presented a substantial innovation by generating stated choice data from a sample of anglers in northeastern Germany that exploited various fish species, including pike. However, different species differ in catch rates and other units of interest (e.g., size dimension), which complicates the standardized estimation of the relative importance of selected attributes across a range of species for angler choice. Beardmore et al. (2013) found a way of tailoring a stated preference discrete choice experiment to a random sample of anglers for which a previous diary survey indicated target species and variation in catch rates and captured sizes to be expected across species by individual respondents. The very same anglers were then confronted in a second survey with a stated choice experiment tailored to their specific target species, where the variation in levels of attributes describing choice option were made species independent by drawing levels for attributes such as catch rates or fish sizes in a standardized fashion across species, thereby varying levels in a comparable way related to species-specifics means and standard deviations for attributes of interest. Thereby, the model generated species-independent estimates of the so-called part worth utilities of different attributes known to be important to anglers, both catch- and non-catch related. To our knowledge this is the most general representation of angler behaviour published so far and hence was chosen for our work. We used these preferences to simulate angler behaviour *in silico*.

In the choice experiment described in detail in Beardmore et al. (2013), randomly selected anglers drawn from fishing license holders in the state of Mecklenburg-Vorpommern (M-V) were presented with a set of hypothetical angling experiences composed of several attributes including target fish species, licence cost, distance to the lake, catch number per trip (catch rate), average and maximum size of catch, number of anglers seen a measure of crowding, minimum-length limit, daily bag limit and stock status. Each attribute was systematically varied to allow estimation of preferences for varying attribute levels. For each choice set, anglers were asked to allocate 10 days among six alternatives: four angling places in the region (i.e., M-V), angling outside the region, and no angling. Besides discrete choice tests, anglers were asked to answer a questionnaire concerning their angling activities during the last twelve months as well as their attitudes towards angling. Random utility theory (McFadden, 1973) assumes that individuals choose one alternative to another to maximize their utility, and the utility of one alternative is a function of its components, i.e., attributes (e.g., expected catch rate) and attribute levels (e.g., different catch rate levels). Based on the observed allocation of days among alternatives, we estimated the part-worth utility (PWU, a measure of importance) for attributes and attribute levels, i.e., the contributions of each attribute and attribute level to the overall utility of the alternative to the angler. We assumed that the PWU for each attribute was a linear function of attribute levels, and estimated the coefficient of the linear function similar to Beardmore et al. (2013). For further details of the choice experiment and its theoretical background, see Beardmore et al. (2013).

### Recreation specialization theory: a framework for understanding angler heterogeneity

Human dimensions researchers have long recognized that the “average angler” does not exist (Aas & Ditton, 1998; Shafer, 1969). In his seminal paper on recreation specialization, Bryan (1977) observed “a continuum of behaviour from the general to the particular, reflected by equipment and skills used in the sport and activity setting preferences” (p. 175) in American trout anglers, concluding that anglers may be grouped into types that share specific values, beliefs, attitudes and behaviours. While conceptualizations of specialization posited that as one gains experience in a recreational activity, one also becomes more emotionally involved or “specialized” (Ditton et al., 1992); however, the notion of clear predictable stages in an angling career being correlated with degree of specialization has been challenged (Scott & Shafer, 2001). That said, specialization is a multidimensional concept (Ditton et al., 1992), with clear correlates related to affective, cognitive and behavioural measures of attachment to the activity (Scott & Shafer, 2001). These measures reflect the degree to which one self-identifies with the activity (Scott & Shafer, 2001), one’s dedication to the values and norms of the social world of angling (Buchanan, 1985; Ditton et al., 1992), one’s level of expertise (Salz & Loomis, 2005), and one’s investment of time, money, and other resources to the activity (Ditton et al., 1992). While these three dimensions form the core of specialization theory, Bryan’s (1977) observation also relies on observations of heterogeneous “activity setting” preferences. Preference can be defined as an evaluative judgment in the sense of liking or disliking an object or outcome (Scherer, 2005). Thus, specialized anglers may also be differentiated from one another by their individual preferences for certain fishing experiences to the exclusion of others. For example, in some fisheries, specialization may be associated with a shift in catch orientation (Anderson et al., 2007; Fedler & Ditton, 1986; Graefe, 1980) from a focus on number of fish towards size of fish; and/or a tendency to release more fish (Bryan, 1977; Salz & Loomis, 2005). In this sense, the concept of specialization may be applied to any segmentation of anglers based on preferences for particular fishing experiences. For example, one may refer to the “fly fisherman” (Bryan, 1977) or “specialized carp angler” (Arlinghaus & Mehner, 2003) as technique or species specialists, or the “trophy angler” (Arterburn et al., 2002) as someone whose behavior is primarily motivated by the outcome of catching a large fish (Fedler & Ditton, 1986, p. 198). While such species-, technique- or outcome-specific preferences may not be fully resolved using the generic model of angler preferences used in this study, specialization still provides a rich conceptual framework for incorporating angler heterogeneity into an examination of social-ecological interactions. We used latent class modelling (see main text) applied to the utility data to identify classes of anglers and investigated whether the angler types followed specialization levels.

One challenge associated with latent class analysis is that the probabilistic nature of the class assignment does not necessarily provide a clear picture of the archetypal member of each class (Beardmore et al., 2013). Examining the angler types identified by latent class analysis through the lens of recreation specialization, however, provided some insights that aided in this regard. For ease of understanding, Table S1 presents a qualitative rating of the relative PWU values among the four angler types for attributes included in the choice experiment, along with other indicators of specialization taken from the surveys, interviews and diaries completed by study participants (for details, see Beardmore et al. 2013; Dorow & Arlinghaus 2011).

Type 1 anglers were the least likely to choose choice alternatives other than fishing in the study region. They were the least averse to paying high license fees, as well as the most accepting of high travel distances to get to fishing destinations within the region. They were the most tolerant of fishing in sight of other anglers, and strict regulations. This group also derived the least utility from the number of fish harvested. The commitment to fishing under less than ideal conditions demonstrated by this group was consistent with their tendency to score highly on a centrality to lifestyle index (measuring the degree to which fishing is a core aspect of their identity and lifestyle, data not shown here), their self-assessment of their fishing skill level, and their financial and travel investments into fishing in the region. These anglers therefore were considered to be highly specialized, fitting the label of “Committed anglers”.

Type 2 anglers and Type 3 anglers represented incremental decreases in specialization, with each of these types more likely to opt out of fishing the last. Of note is the apparently increasing importance of catch outcomes among these groups, indicating that perceptions of high fishing quality are necessary to overcome the propensity to pursue non-fishing activities. Type 2 and Type 3 anglers were therefore considered to exhibit moderate and low specialization levels, respectively, fitting the labels of active and casual anglers.

While the first three angler types represent a specialization continuum from committed to casual in their preferences and commitment to angling in the study region, Type 4 anglers presented a different breed. In their choice responses, Type 4 anglers demonstrated a strong preference for fishing outside the region, showed a medium aversion to license costs and travel within the region. They derived higher utility from larger fish. They considered themselves to be more skilled on average than did the other groups, but were similar to Type 2 in their centrality to lifestyle. On average they tended to travel farthest to fish, while paying less than other anglers for their regional licenses. On the other hand, they had the highest average investment in fishing equipment. On the whole, Type 4 anglers appeared less invested in fishing freshwaters in Mecklenburg-Vorpommern; however, their commitment to fishing extended beyond the borders of the state with substantial investments in time and money to pursue their fishing activities. Consequently, one should consider these anglers as highly specialized (similar to Type 1 anglers) but with a greater emphasis on fishing elsewhere than in the study region.

### Calibration of the mechanistic angler model

The original choice model presented the levels of some attributes (in particular catch rates, the size of fish captured and the angler numbers seen while fishing) in a standardized and personalized fashion to remove scale and units issues different among species (Beardmore et al. 2013). This was done by varying the levels of the mentioned attributes around a species specific distribution in units of SD so that choice sets presented to respondents for species A and species B varied in the same fashion along species-specific characteristics (e.g., the same SD change in length of fish captured when a pike scenario was evaluated compared to a perch scenario, for example). To find means and SDs for the attributes Fish number, Maximum size, and Angler seen in our simulated virtual landscape and allow the calibration of Beardmore et al.’s (2013) model, initial simulations were run assuming that anglers are distributed across the lakes to achieve the maximum sustainable yield (MSY) of any one population. The resulting distribution of the three attribute levels at equilibrium across lakes were used to define the expected variation in the virtual landscape at optimal conditions and to compute means and SDs so that variation in catch rate, size and crowding all exerted effect on utility, and hence on lake choice.

For application of the choice model to our landscape, we modified the originally estimated linear PWU function for the utility effect of catch rates (fish numbers, Table S1). This was done because during preliminary simulations using the original functions given by Beardmore et al. (2013), we found that anglers also visited lakes where catch rates are zero. This unreasonable outcome arouse from the fact that the original stated preference choice experiment did not include extremely low catch levels by design. Moreover, the extreme nonlinearities of the PWU function for fish catch reported in subsequent work by Arlinghaus et al. (2014) (i.e., infinitely low utility of zero catch and marginal diminishing returns as catch rates close some threshold level of one to two fish per day) could not be approximated by the original linear function fitted through five catch levels in the experiment by Beardmore et al. (2013). To avoid systematically overestimating the number of anglers at lakes present at even extremely low catch rates, we re-fitted logarithmic functions

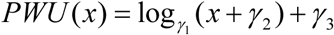

through the PWU values predicted by the original functions at five levels of expected catch rates (*x*), that is, *μ* − 2.63*σ*, *μ* − 0.5*σ*, *μ*, *μ* + 1.0*σ*, and *μ* + 3.76*σ*, where *μ* and *σ* represent the mean and SD for catch rates at MSY, respectively. The first and last values correspond to actual catch rates that are zero and the maximum number of catch rates per angler possible in the region, respectively. Note that we a standardized catch rate before calculating PWU, so the mean of the standardized catch rate is zero and the absolutely zero value for catch rate is negative on the standardized curve (Fig. S1). The PWU at the point of actual zero catch rate was determined to achieve a low probability of fishing of 6.3% when PWUs of all other attributes are zero. The probability value (6.3%) was chosen corresponding with angler diary data from anglers in M-V; it corresponds to the average percentage of trips taken by anglers who had average daily catch rates of zero. The modified functions are shown in Fig. S1, and the values of parameters *γ*_1_, *γ*_2_, and *γ*_3_ are reported in Table S1. The functional form agreed with the diminishing marginal return of utility of catch rate expected from economic theory and reported for German anglers elsewhere (Arlinghaus et al. 2014). In Fig. S1 you can also see variation in angler types in how utility of catch rate changes with increasing catch.

We also modified regulation-related attributes in Beardmore et al. (2013). We combined the original attributes “Minimum-size limit” and “Daily bag limit” and created a single attribute “Regulations”, and estimated parameter values of the new PWU function.

---

**Table S1.** Relative utility values across angler types and indicator variables associated with for classification by recreation specialization assessed from the choice model (see main text) and a qualitative assessment of differences among anglers in additional variables (cognitive and affective as well as behaviour) taken from the survey data (for details see Beardmore et al., 2013).

|  | **Type 1** | **Type 2** | **Type 3** | **Type 4** |
| --- | --- | --- | --- | --- |
| **Attribute** |  |  |  |  |
| Propensity to Fish | High (in region) | Medium | Low | High (Elsewhere) |
| Cost aversion | Very Low | Low | High | Medium |
| Travel aversion | Low | High | Very High | Medium |
| Utility from fish harvested | Low | Medium | High | Medium |
| Utility from max. size | Medium | Medium | Medium | High |
| Congestion aversion | Very Low | Medium | Low | Medium |
| Overfishing aversion | High | High | Medium | High |
| Regulation aversion | Low | Medium | High | Medium |
| **Cognitive and Affective commitment** |  |  |  |  |
| Centrality to lifestyle (affective) | High | Medium | Low | Medium |
| Self-rated angling skill (cognitive) | Medium-High | Medium | Low | High |
| **Behavioral commitment** |  |  |  |  |
| Average travel distance | High | Medium | Low | Very high |
| License expenditures in MV | High | High | Medium | Low |
| Average trips targeting pike (per year) | 4.3 | 4.3 | 3.6 | 3.3 |
| Equipment value (Euro) | 1520 | 1120 | 913 | 1834 |
| **Specialization label** | **Committed (in region)** | **Active** | **Casual** | **Committed (elsewhere)** |

**Fig. S1.**
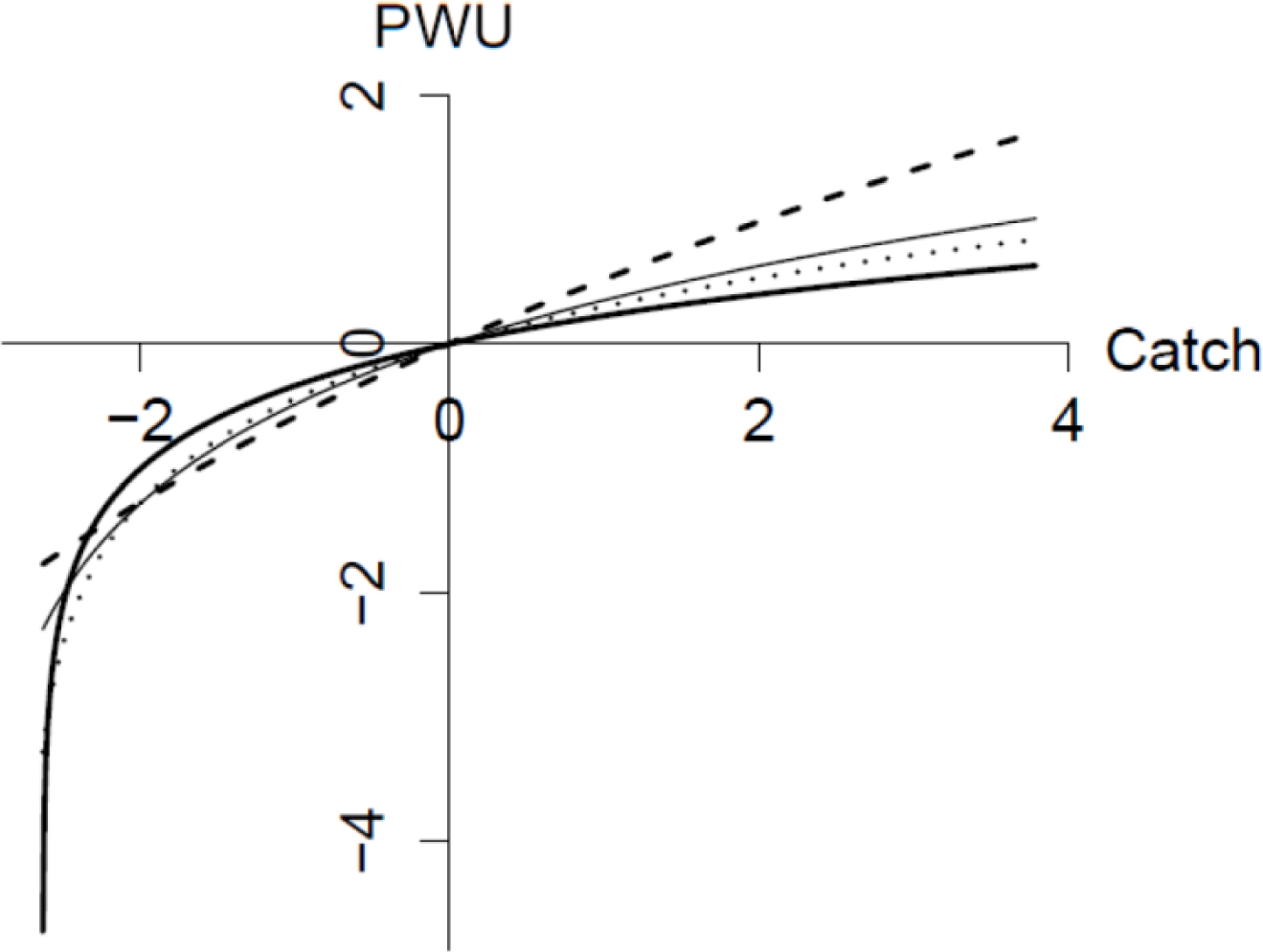
Modified part worth utility (PWU) functions for standardized catch rate. The smallest value of the standardized catch rate corresponds to zero catch. Thick, dotted, dashed and thin lines correspond to the type 1, 2, 3, and 4 angler classes (Table S1), respectively.

## Appendix II

**Figure S2.**
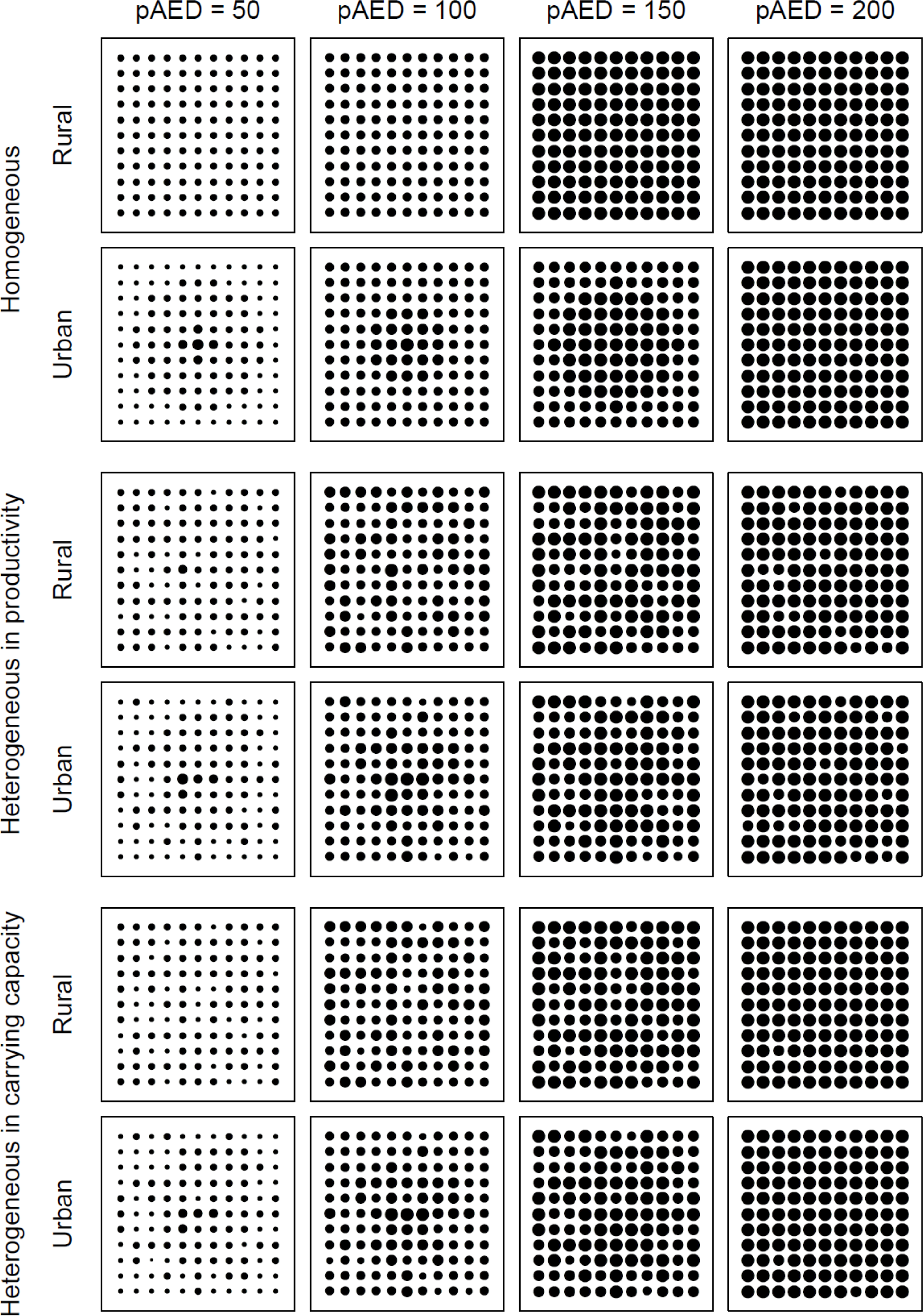
An example of the distribution of lake-specific angling effort in the homogeneous and heterogeneous landscapes in the absence of harvest regulations in the rural and urban landscape. Lakes are identical in the homogeneous landscape, while lakes differ in their productivity or carrying capacity in the heterogeneous landscape. The annual angling effort densities are: <30, <60, <90, <120, <150, and ≥150 [h ha^-1^].

**Figure S3.**
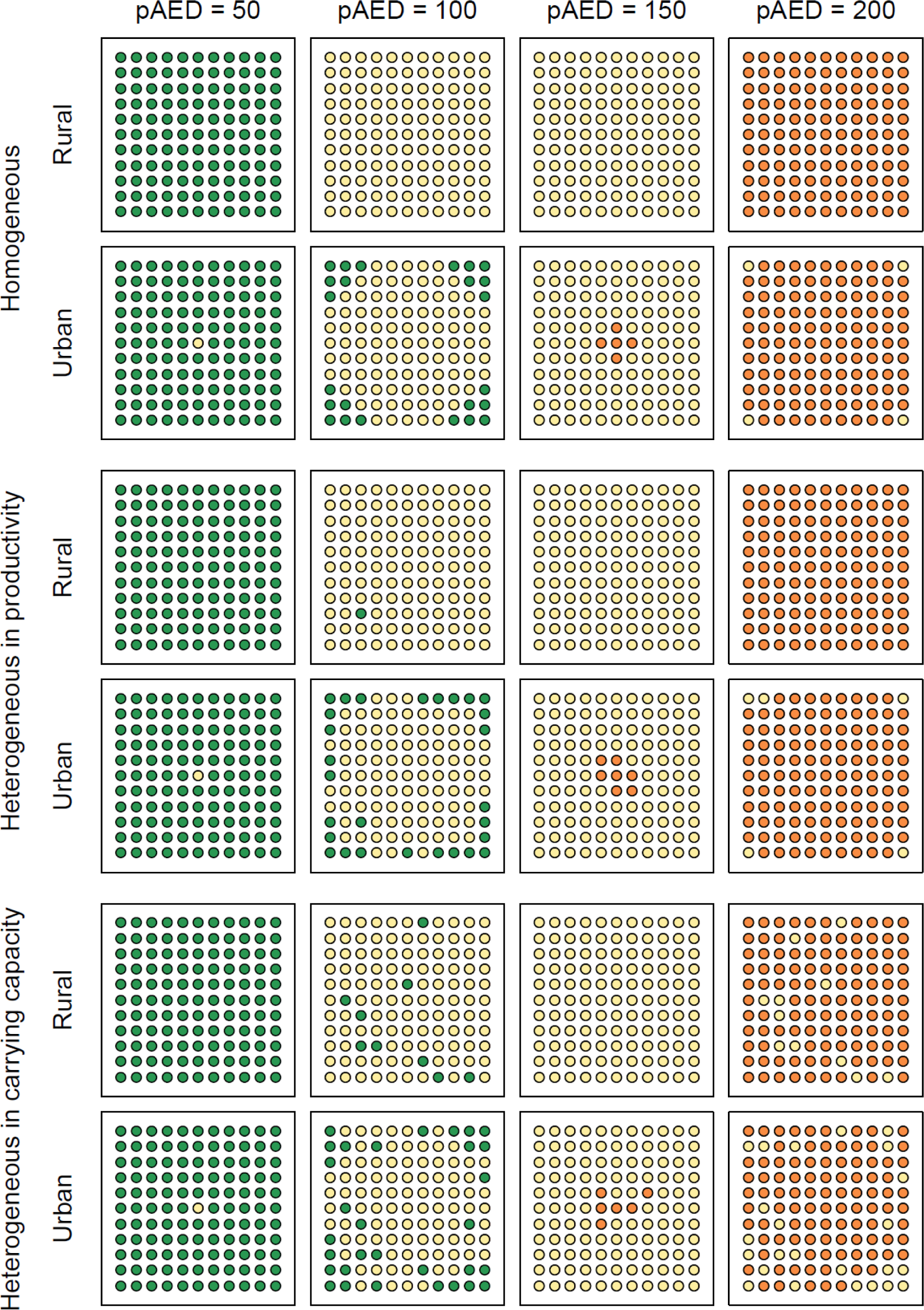
An example of the spatial pattern of exploitation in the homogeneous and heterogeneous landscapes in the absence of harvest regulations in the rural and urban landscapes. Lakes are identical in the homogeneous landscape, while lakes differ in their productivity or carrying capacity in the heterogeneous landscape. Lakes are categorized based on their relative spawning stock biomass (SSB) to their pristine SSB (SSB/SSB_0_). Green: healthy (0.35 or higher), yellow: overfished (between 0.35 and 0.10), red: collapsed (less than 0.10). pAED is potential angling effort density [h ha^-1^].

**Figure S4.**
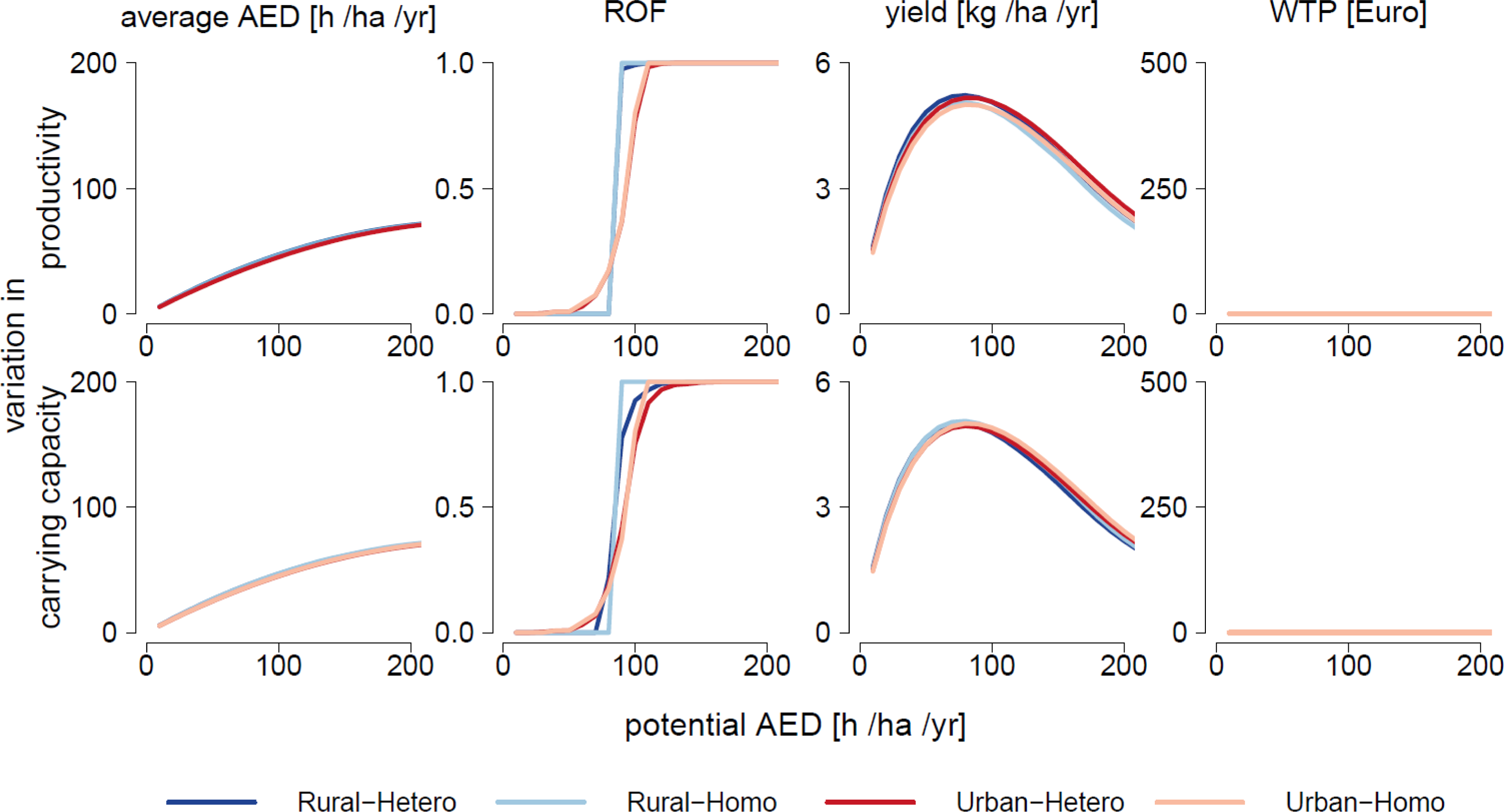
Comparison between the homogeneous (Homo) and heterogeneous (Hetero) landscapes. Lakes are identical in the homogeneous landscape, while lakes vary in their productivity (top) or carrying capacity (bottom) in the heterogeneous landscape. Regional outcomes in terms of average lake-specific angling effort, degree of overexploitation of lakes (ROF = recruitment overfished stocks), biomass yield, and angler welfare as represented by average willingness-to-pay (WTP) per year in the rural and urban landscapes in the absence of harvest regulations are shown.

**Figure S5.**
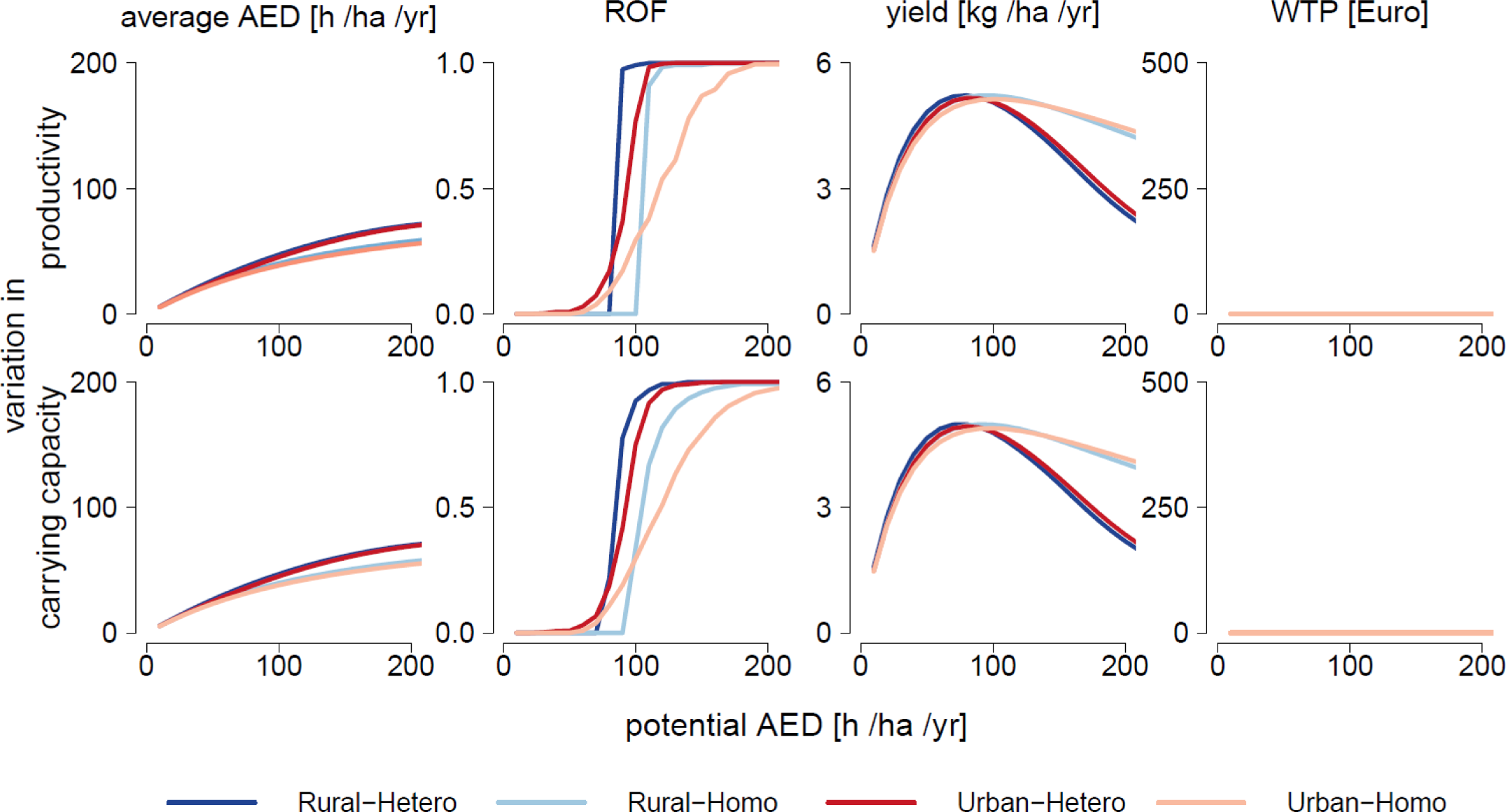
Comparison between the 4-class heterogeneous (Hetero) and 1-class homogeneous (Homo) angler models in the rural and urban landscapes. Regional outcomes in angling effort, overexploitation of lakes (ROF = recruitment overfished lakes), biomass yield and angler welfare as represented by average willingness-to-pay (WTP) per year in the absence of harvest regulations are shown. Lakes vary in their productivity (top) or carrying capacity (bottom).

**Figure S6.**
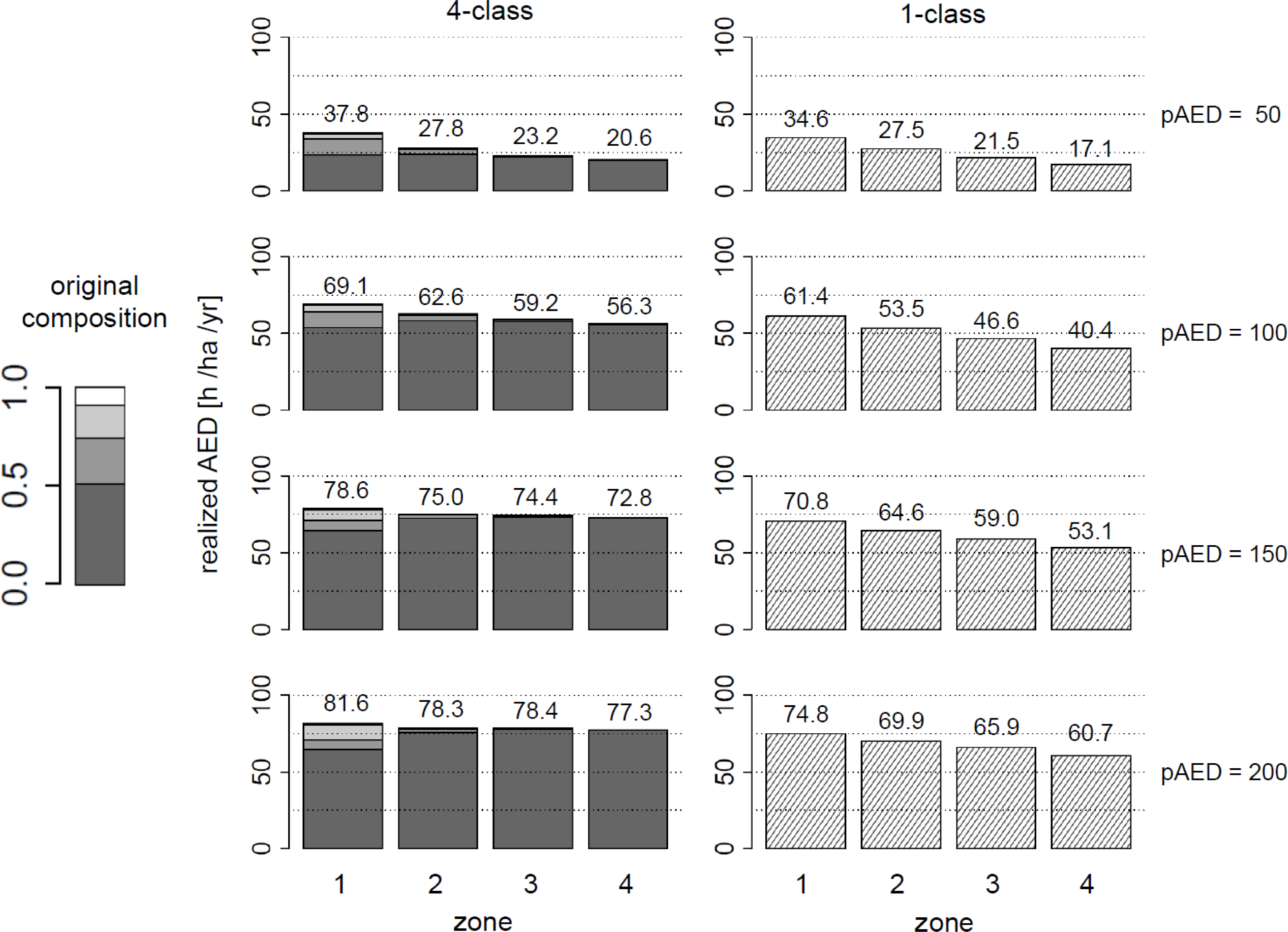
Proportions of each angler class within the realised angling effort density (AED, angling-h ha^-1^) in the urban case in the absence of harvest regulations. Lakes vary in their productivity. Lakes are categorized by the distance from the metropolis: Zone 1 (<28 km), 2 (<56 km) 3 (<84 km) and 4 (≥84 km). The original proportion of the angler classes is shown on the left.

**Figure S7.**
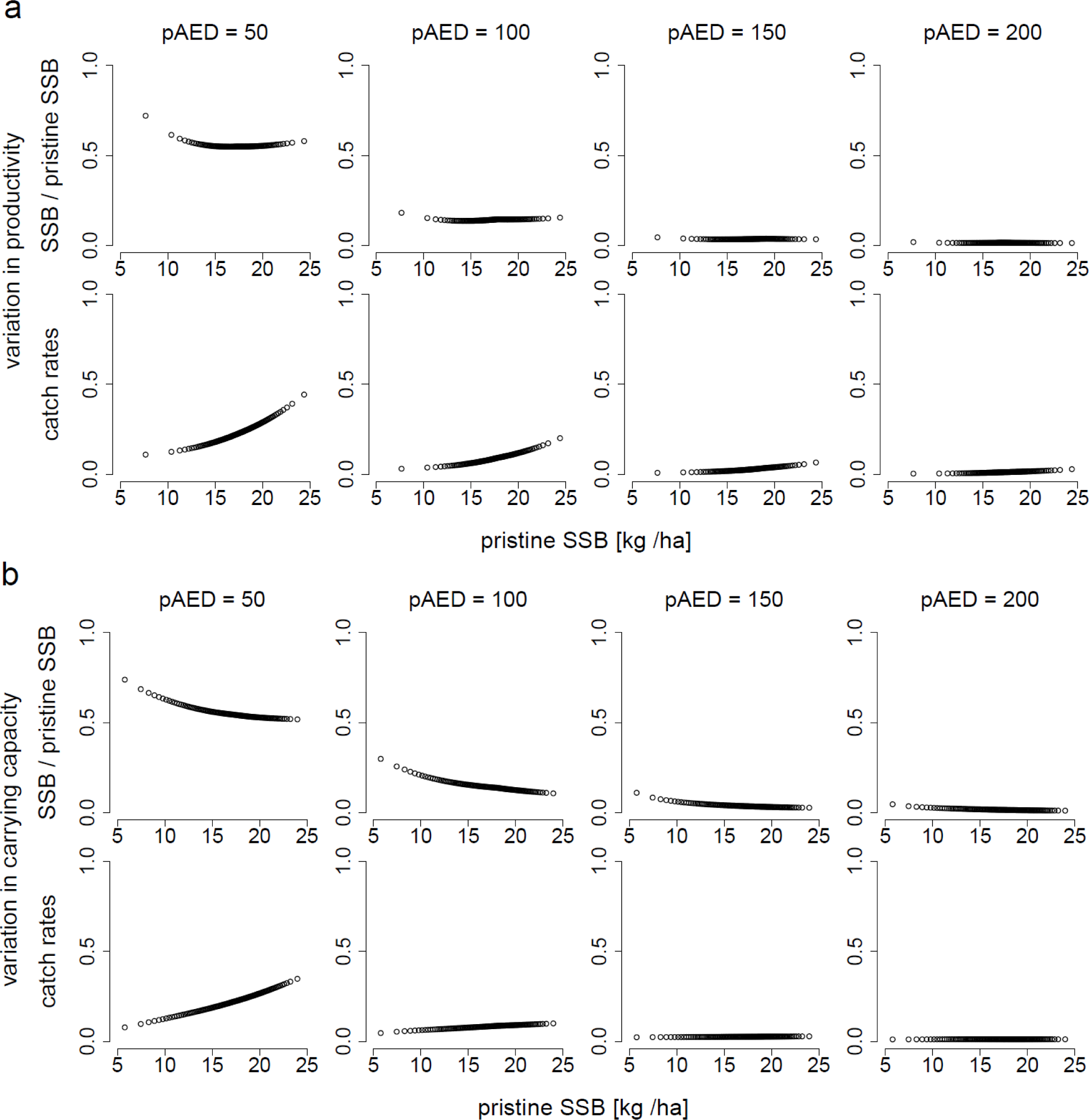
Relationship between a lake’s intrinsic quality (pristine SSB = SSB_0_) and the degree of exploitation (represented by SSB/SSB_0_) and average angler catch rates (pike per hour) at equilibrium in the absence of harvest regulations in a rural landscape. Each lake is represented by a circle. Variation among lakes in their pristine SSB arises either from variation in their productivity (a) or carrying capacity (b). From the left to the right: potential pAED (annual angling effort density) = 50, 100, 150, or 200 [h ha^-1^].

**Figure S8.**
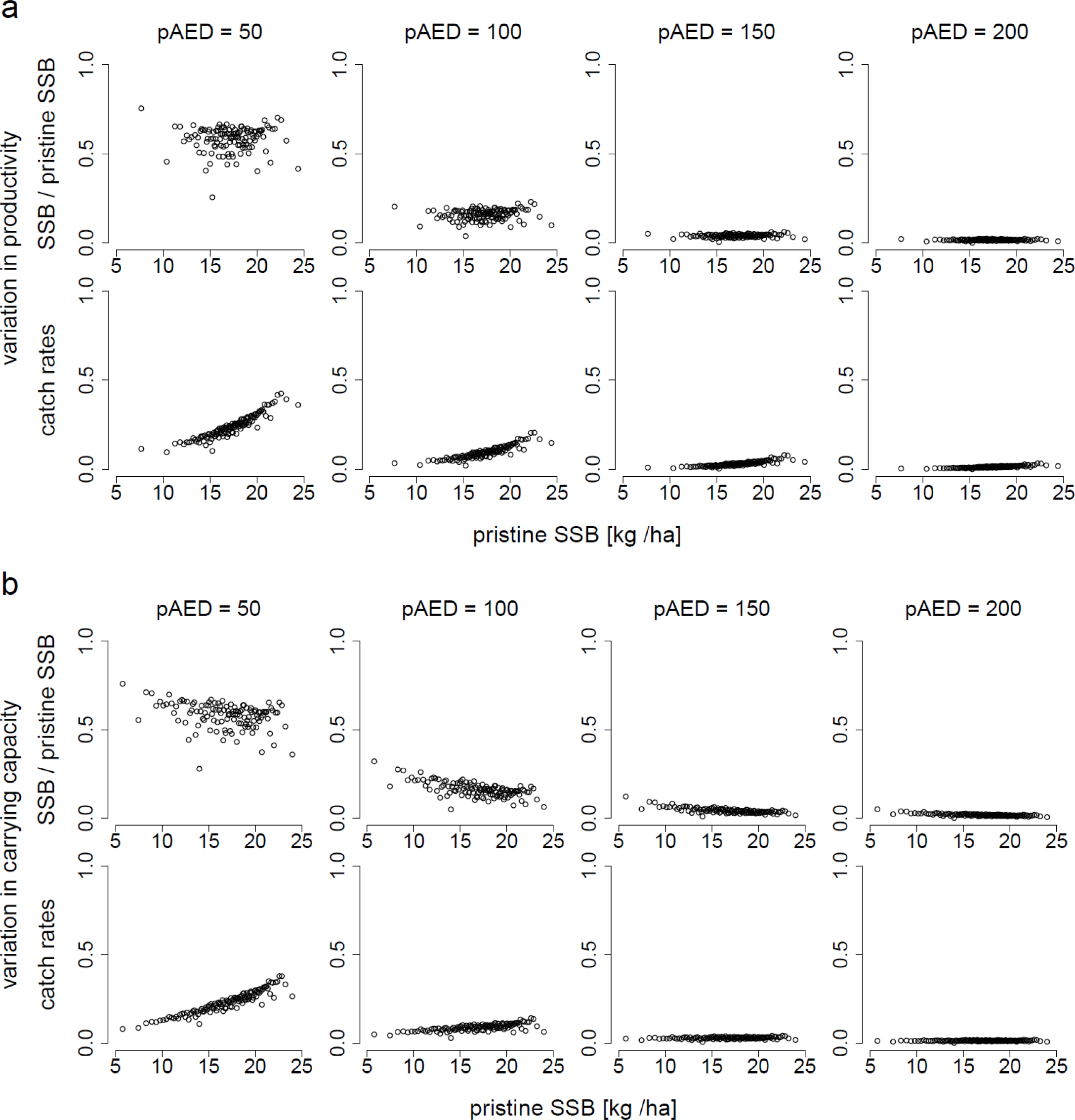
Relationship between a lake’s intrinsic quality (pristine SSB = SSB_0_) and the degree of exploitation (represented by SSB/SSB_0_) and average angler catch rates (pike per hour) at equilibrium in the absence of harvest regulations in an urban landscape. Each lake is represented by a circle. Variation among lakes in their pristine SSB arises either from variation in their productivity (a) or carrying capacity (b). From the left to the right: potential pAED (annual angling effort density) = 50, 100, 150, or 200 [h ha^-1^].

**Figure S9.**
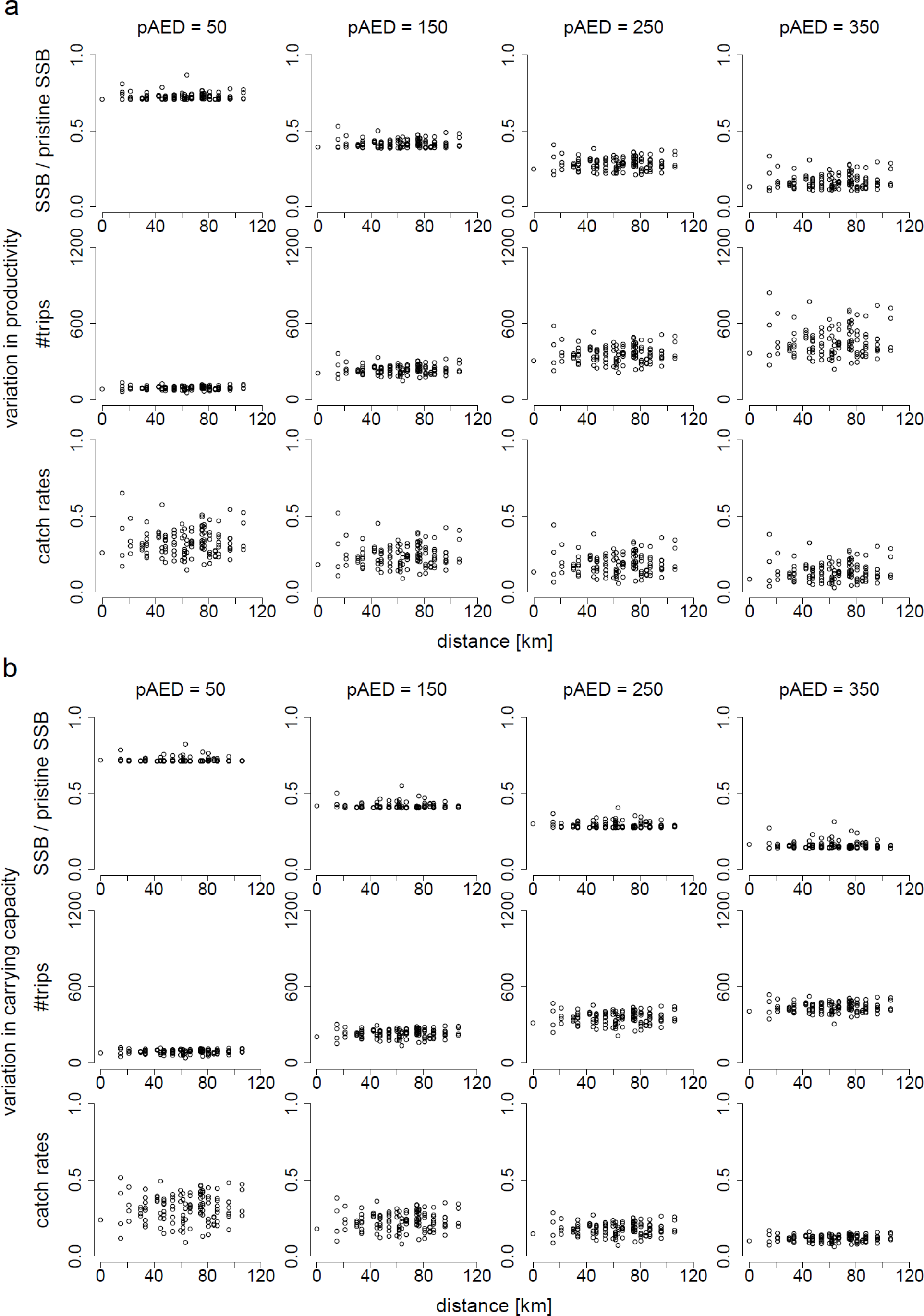
Relationship between the distance from the central lake and the degree of exploitation (represented by SSB/SSB_0_), the number of trips taken per year to each lake, and average angler catch rates (pike per hour) at equilibrium in the absence of harvest regulations in a rural landscape. Each lake is represented by a circle. Variation among lakes in their pristine SSB arises either from variation in their productivity (a) or carrying capacity (b). From the left to the right: potential AED (annual angling effort density) = 50, 100, 150, and 200 [h ha^-1^].

**Figure S10.**
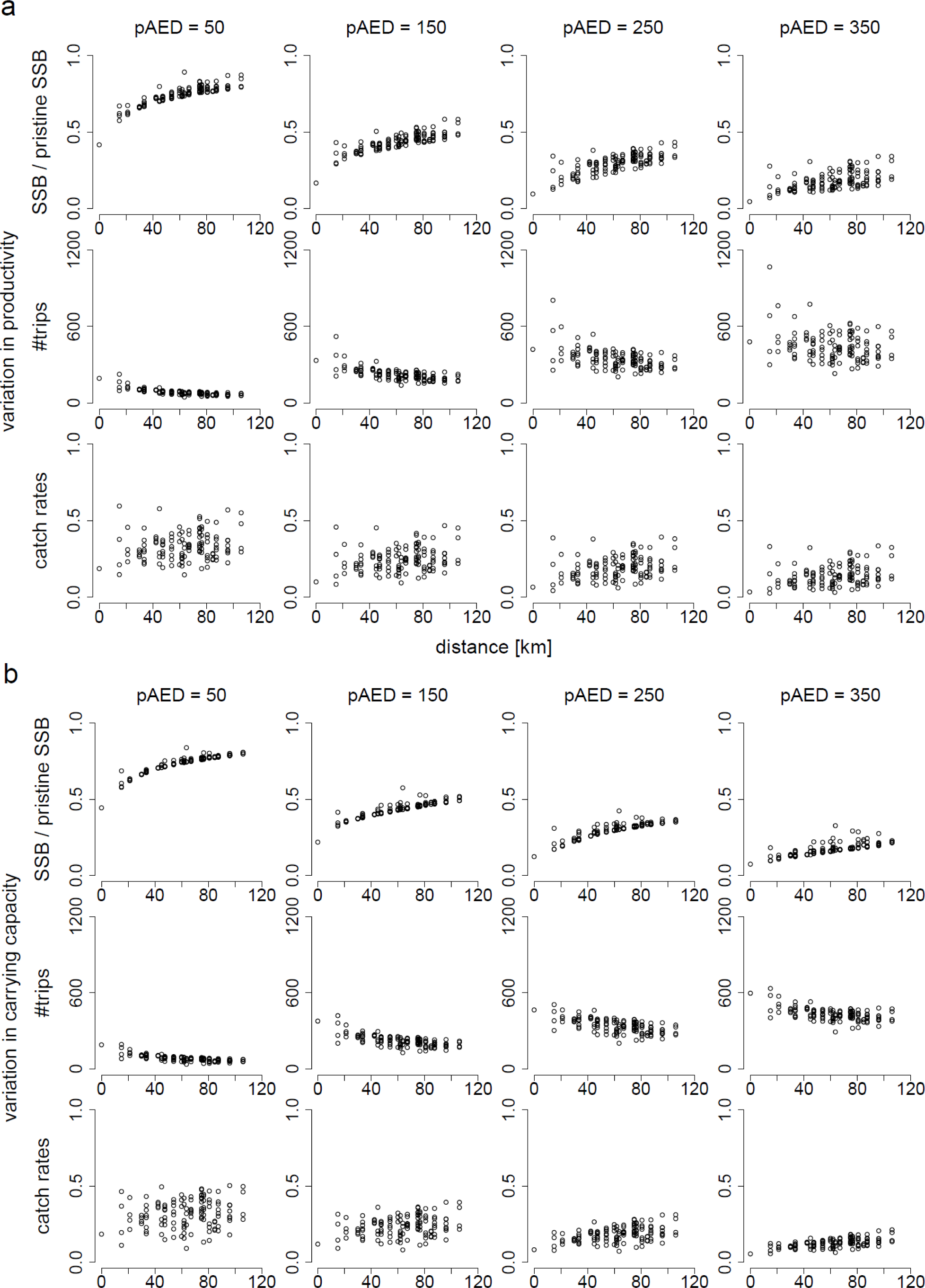
Relationship between the distance from the central lake and the degree of exploitation (represented by SSB/SSB_0_), the number of trips taken per year to each lake, and average angler catch rates (pike per hour) at equilibrium in the absence of harvest regulations in an urban landscape. Each lake is represented by a circle. Variation among lakes in their pristine SSB arises either from variation in their productivity (a) or carrying capacity (b). From the left to the right: potential AED (annual angling effort density) = 50, 100, 150, and 200 [h ha-^1^].

**Figure S11.**
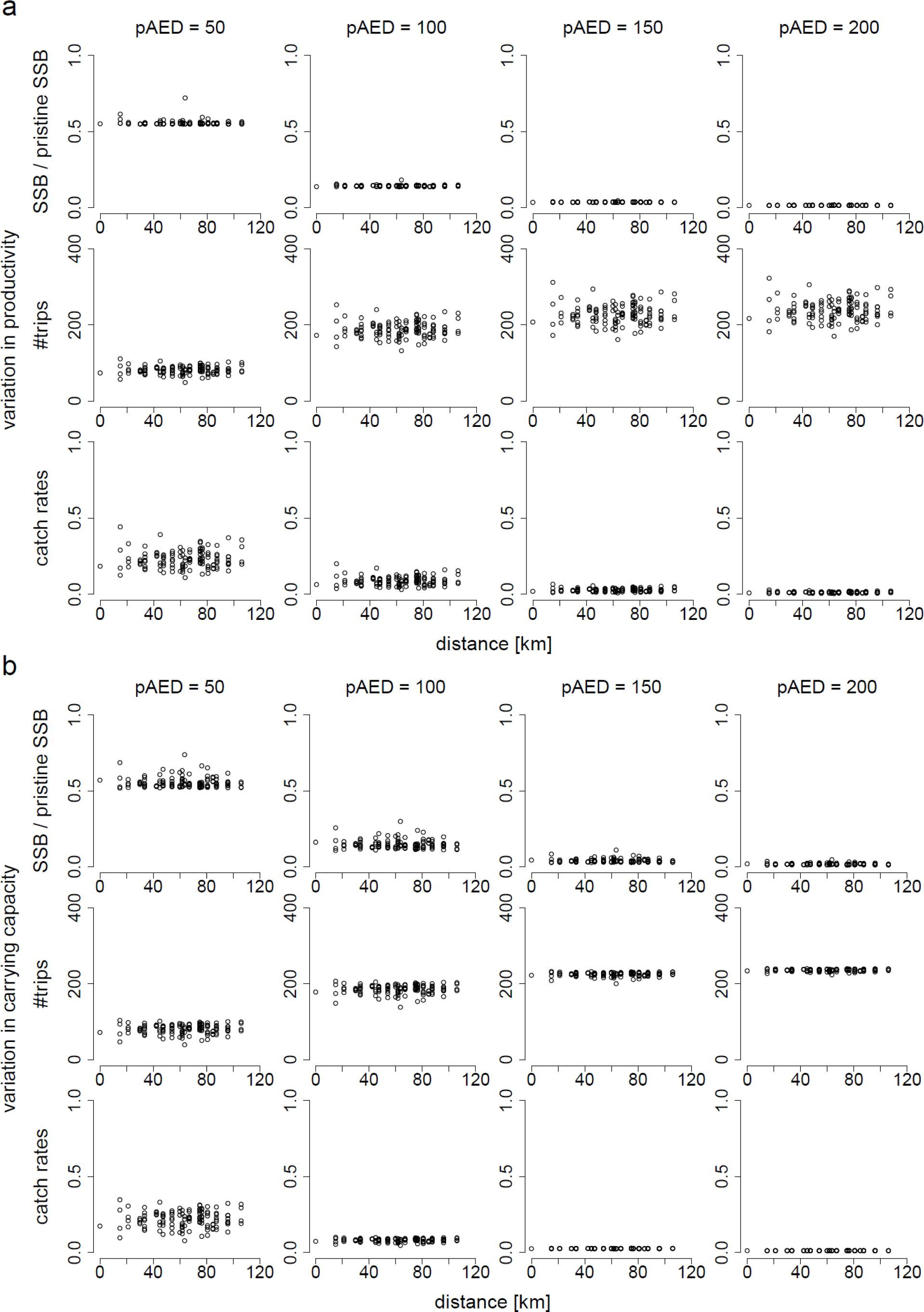
Relationship between the distance from the central lake and the degree of exploitation (represented by SSB/SSB_0_), the number of trips taken per year to each lake, and average angler catch rates (pike per hour) at equilibrium with the presence of the one-size-fits all harvest regulation in a rural landscape. Each lake is represented by a circle. Variation among lakes in their pristine SSB arises either from variation in their productivity (a) or carrying capacity (b). From the left to the right: potential AED (annual angling effort density) = 50, 150, 250, and 350 [h ha^-1^].

**Figure S12.**
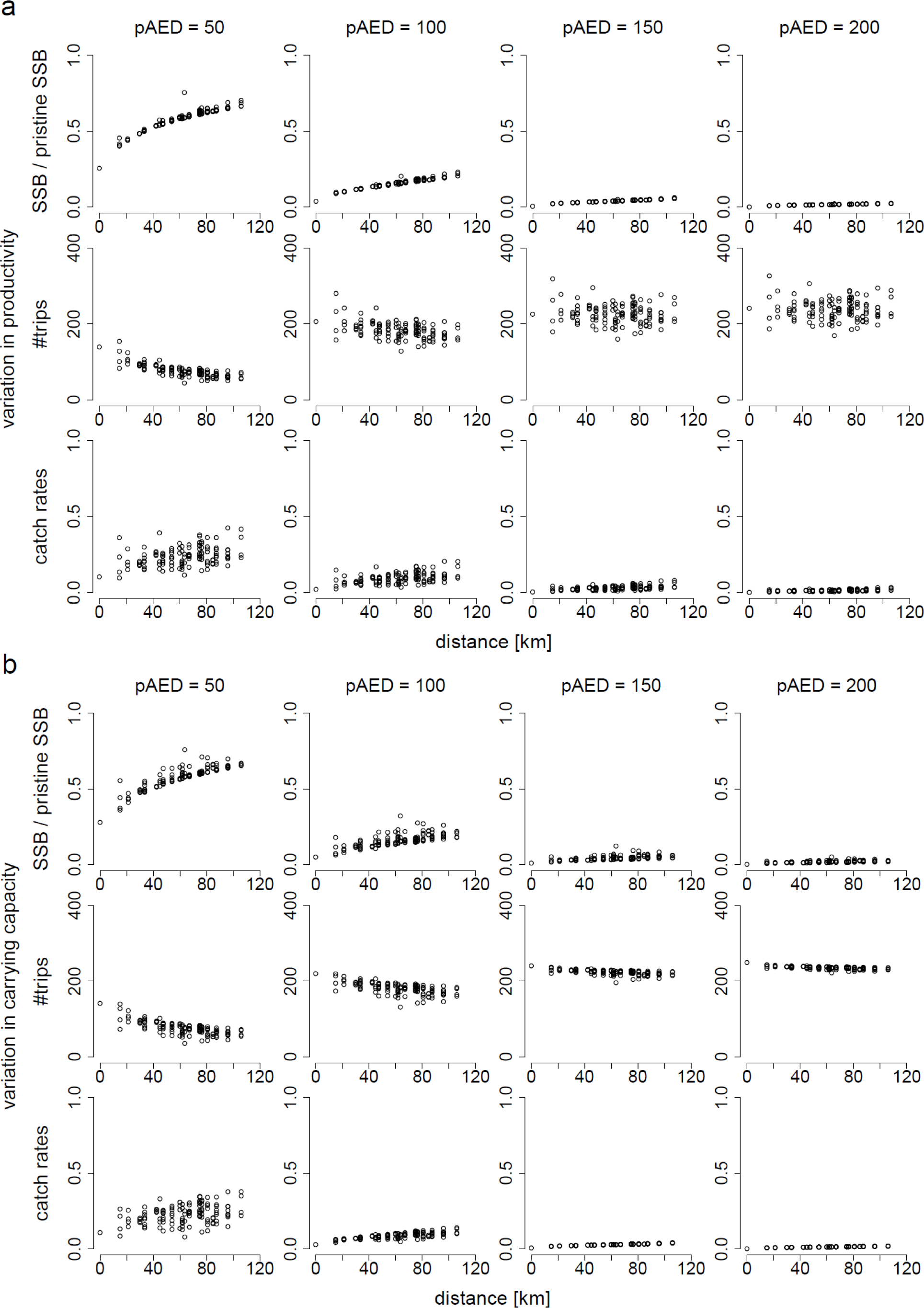
Relationship between the distance from the central lake and the degree of exploitation (represented by SSB/SSB_0_), the number of trips taken per year to each lake, and average angler catch rates (pike per hour) at equilibrium with the presence of the one-size-fits all harvest regulation in an urban landscape. Each lake is represented by a circle. Variation among lakes in their pristine SSB arises either from variation in their productivity (a) or carrying capacity (b). From the left to the right: potential AED (annual angling effort density) = 50, 150, 250, and 350 [h ha^-1^].

## References

Abbott, J. K., & Fenichel, E. P. (2013). Anticipating adaptation: a mechanistic approach for linking policy and stock status to recreational angler behavior. Canadian Journal of Fisheries and Aquatic Sciences, 70, 1190–1208. doi:10.1139/cjfas-2012-0517

Allen, M. S., Brown, P., Douglas, J., Fulton, W., & Catalano, M. (2009). An assessment of recreational fishery harvest policies for Murray cod in southeast Australia. Fisheries Research, 95, 260–267. doi:10.1016/j.fishres.2008.09.028

Allen, M. S., Ahrens, R. N. M., Hansen, M. J., & Arlinghaus, R. (2013). Dynamic angling effort influences the value of minimum-length limits to prevent recruitment overfishing. Fisheries Management and Ecology, 20, 247–257. doi:10.1111/j.1365-2400.2012.00871.x

Anderson, L. G. (1993). Toward a complete economic theory of the utilization and management of recreational fisheries. Journal of Environmental Economics and Management, 24, 272–295. doi:10.1006/jeem.1993.1018

Arlinghaus, R., & Mehner, T. (2005). Determinants of management preferences of recreational anglers in Germany: habitat management versus fish stocking. Limnologica, 35, 2–17. doi: 10.1016/j.limno.2004.10.001

Arlinghaus, R., Matsumura, S., & Dieckmann, U. (2009). Quantifying selection differentials caused by recreational fishing: development of modeling framework and application to reproductive investment in pike (*Esox lucius*). Evolutionary Applications, 2, 335–355. doi:10.1111/j.1752-4571.2009.00081.x

Arlinghaus, R., Matsumura, S., & Dieckmann, U. (2010). The conservation and fishery benefits of protecting large pike (*Esox lucius* L.) by harvest regulations in recreational fishing. Biological Conservation, 143, 1444–1459. doi:10.1016/j.biocon.2010.03.020

Arlinghaus, R., Cooke, S. J., & Potts, W. (2013). Towards resilient recreational fisheries on a global scale through improved understanding of fish and fisher behaviour. Fisheries Management and Ecology, 20, 91–98. doi:10.1111/fme.12027

Arlinghaus, R., Beardmore B., Riepe, C., Meyerhoff, J., & Pagel, T. (2014). Species-specific preferences of German recreational anglers for freshwater fishing experiences, with emphasis on the intrinsic utilities of fish stocking and wild fishes. Journal of Fish Biology, 85, 1843–1867. doi:10.1111/jfb.12546

Arlinghaus, R., Cyrus, E.-M., Eschbach, E., Fujitani, M., Hühn, D., Johnston, F., Pagel, T., & Riepe, C. (2015). Hand in Hand für eine nachhaltige Angelfischerei: Ergebnisse und Empfehlungen aus fünf Jahren praxisorientierter Forschung zu Fischbesatz und seinen Alternativen. Berichte des IGB, Heft 28.

Arlinghaus, R., Lorenzen, K., Johnson, B. M., Cooke, S. J., & Cowx, I. J. (2016). Management of freshwater fisheries. In J. F. Craig (Ed.), Freshwater fisheries ecology (pp. 557–579). Chichester, UK: John Wiley & Sons, Ltd.

Arlinghaus, R., Alós, J., Beardmore, B., Daedlow, K., Dorow, M., Fujitani, M., Hühn, D., Haider, W., Hunt, L. M., Johnson, B. M., Johnston, F., Klefoth, T., Matsumura, S., Monk, C., Pagel, T., Post, J. R., Rapp, T., Riepe, C., Ward, H., & Wolter, C. (2017). Understanding and managing freshwater recreational fisheries as complex adaptive social-ecological systems. Reviews in Fisheries Science and Aquaculture, 25, 1–41. doi:10.1080/23308249.2016.1209160

Askey, P. J., Richards, S. A., Post, J. R., & Parkinson, E. A. (2006). Linking angling catch rates and fish learning under catch-and-release regulations. North American Journal of Fisheries Management, 26, 1020–1029. doi:10.1577/M06-035.1

Askey, P. J., Parkinson, E. A., & Post, J. R. (2013). Linking fish and angler dynamics to assess stocking strategies for hatchery-dependent, open-access recreational fisheries. North American Journal of Fisheries Management, 33, 557–568. doi:10.1080/02755947.2013.785996

Beardmore, B., Haider, W., Hunt, L. M., & Arlinghaus, R. (2011). The importance of trip context for determining primary angler motivations: Are more specialized anglers more catch-oriented than previously believed? North American Journal of Fisheries Management, 31, 861–879. doi:10.1080/02755947.2011.629855

Beardmore, B., Haider, W., Hunt, L. M., & Arlinghaus, R. (2013). Evaluating the ability of specialization indicators to explain fishing preferences. Leisure Science, 35, 273–292. doi: 10.1080/01490400.2013.780539

Bryan, H. (1977). Leisure value systems and recreational specialization: the case of trout fishermen. Journal of Leisure Research, 9, 174–187. doi:10.1177/004728757801600352

Camp, E. V., van Poorten, B. T., & Walters, C. J. (2015). Evaluating short openings as a management tool to maximize catch related utility in catch-and-release fisheries. North American Journal of Fisheries Management, 35, 1106–1120. doi:10.1080/02755947.2015.1083495

Camp, E. V., Larkin, S. L., Ahrensa, R. N. M., & Lorenzen, K. (2017). Trade-offs between socioeconomic and conservation management objectives in stock enhancement of marine recreational fisheries. Fisheries Research, 186, 446–459. doi:10.1016/j.fishres.2016.05.031

Carpenter, S. R., & Brock, W. A. (2004). Spatial complexity, resilience and policy diversity: fishing on lake-rich landscapes. Ecology and Society, 9, 8.

Carpenter, S. R., Munoz-Del-Rio, A., Newman, S., Rasmussen, P. W., & Johnson, B. M. (1994). Interactions of anglers and walleyes in Escanaba Lake, Wisconsin. Ecological Applications, 4, 822–832. doi:10.2307/1942011

Carpenter, S. R., Brock, W. A., Folke, C., van Nese, E. H., & Scheffer, M. (2015). Allowing variance may enlarge the safe operating space for exploited ecosystems. Proceedings of the National Academy of Sciences of the United States of America, 112, 14384–14389. doi:10.1073/pnas.1511804112

Cole, R. A., & Ward, F. A. (1994). Optimum fisheries management policy: angler opportunity versus angler benefit. North American Journal of Fisheries Management, 14, 22–33. doi:10.1577/1548-8675(1994)014<0022:OFMPAO>2.3.CO;2

Cox S. P., & Walters, C. (2002). Maintaining quality in recreational fisheries: how success breeds failure in management of open-access sport fisheries. In T. J. Pitcher & C. E. Hollingworth (Eds.), Recreational fisheries: Ecological, economic and social evaluation (pp. 107–119). Oxford, UK: Blackwell Publishing.

Craig, J. F., & Kipling, C. (1983). Reproduction effort versus the environment; case histories of Windermere perch, *Perca fluviatilis* L., and pike, *Esox lucius* L. Journal of Fish Biology, 22, 713–727. doi:10.1111/j.1095-8649.1983.tb04231.x

Crane, D. P., Miller, L. M., Diana, J. S., Casselman, J. M., Farrell, J. M., Kapuscinski, K. L., & Nohner, J. K. (2015). Muskellunge and northern pike ecology and management: important issues and research needs. Fisheries, 40, 258–267. doi:10.1080/03632415.2015.1038382

Daedlow, K., Beard, T. D. Jr., & Arlinghaus. R. (2011). A property rights-based view on management of inland recreational fisheries: contrasting common and public fishing rights regimes in Germany and the U.S.A. In T. D. Beard Jr., R. Arlinghaus, & S. Sutton (Eds.), Proceedings of the 5th World Recreational Fishing Conference (pp. 13–18), American Fisheries Society Symposium 75. Bethesda, MD: American Fisheries Society.

Deroba, J. J., & Bence, J. R. (2008). A review of harvest policies: Understanding relative performance of control rules. Fisheries Research, 94, 210–223. doi:10.1016/jfishres.2008.01.003

Diana, J. S. (1983). Growth, maturation, and production of northern pike in three Michigan lakes. Transactions of the American Fisheries Society, 112, 38–46. doi:10.1577/1548-8659(1983)112<38:GMAP0N>2.0.C0;2

Ditton, R. B, Loomis, D. K., & Choi, S. (1992). Recreation specialization: Reconceptualization from a social worlds perspective. Journal of Leisure Research, 24, 33–51.

Dorow, M., Beardmore, B., Haider, W., & Arlinghaus, R. (2010). Winners and losers of conservation policies for European eel, *Anguilla anguilla:* an economic welfare analysis for differently specialised eel anglers. Fisheries Management and Ecology, 17, 106–125. doi:10.1111/j.1365-2400.2009.00674.x

Edline, E., Carlson, S. M., Stige, L. C., Winfield, I. J., Fletcher, J. M., James, J. B., & Haugen, T. O. (2007). Trait changes in a harvested population are driven by a dynamic tug-of-war between natural and harvest selection. Proceedings of the National Academy of Sciences of the United States of America, 104, 15799–15804. doi:10.1073/pnas.0705908104

Fayram, A. H., Schenborn, D. A., Hennessy, J. M., Nate, N. A., & Schmalz, P. J. (2009). Exploring the conflict between broad scale and local inland fisheries management: the risks to agency credibility. Fisheries, 34, 232–236. doi:10.1577/1548-8446-34.5.232

Fenichel, E. P., & Abbott, J. K. (2014). Heterogeneity and the fragility of the first best: putting the “micro” in bioeconomic models of recreational resources. Resources and Energy Economics, 36, 351–369. doi:10.1016/j.reseneeco.2014.01.002

Fenichel E. P., Abbott, J. K., & Huang, B. (2013a). Modelling angler behaviour as a part of the management system: synthesizing a multi-disciplinary literature. Fish and Fisheries, 14, 137–157. doi:10.1111/j.1467-2979.2012.00456.x

Fenichel, E. P., Gentner, B., & Arlinghaus, R. (2013b). Normative considerations for recreational fishery management: a bioeconomic framework for linking positive science and normative fisheries policy decisions. Fisheries Management and Ecology, 20, 223–233. doi:10.1111/j.1365-2400.2012.00869.x

Franklin, D. R., & Smith, L. L. Jr. (1963). Early life history of the northern pike *(Esox lucius* L.) with special reference to the factors influencing the numerical strength of year classes. Transactions of the American Fisheries Society, 92, 91–110. doi:10.1577/1548-8659(1963)92[91:ELHOTN]2.0.CO;2

Fretwell, S. D., & Lucas, H. L. Jr. (1970). On territorial behaviour and other factors influencing habitat distribution in birds. I. Theoretical development. Acta Biotheoretica, 19, 16–36. doi:10.1007/BF01601954

Gale, R. P. (1987). Resource miracles and rising expectations: a challenge to fishery managers. Fisheries, 12, 8–13. doi:10.1577/1548-8446(1987)012<0008:RMAREA>2.0.CO;2

Hahn, J. (1991). Angler specialization: measurement of a key sociological concept and implications for fisheries management decisions. In D. Guthrie, J. M. Hoenig, M. Holliday, C. M. Jones, M. J. Mills, S. A. Moberly, K. H. Pollack, & D. R. Talhelm (Eds.), Creel and angler surveys in fisheries management: Proceedings of the International Symposium and Workshop on Creel and Angler Surveys in Fisheries Management, held at Houston, Texas, USA, 26–31 March 1990 (pp. 380–389). Bethesda, MD: American Fisheries Society.

Hahnemann, W. M. (1984). Welfare evaluation in contingent valuation experiments with discrete responses. American Journal of Agricultural Economics, 66, 332–341. doi:10.2307/1240800

Haugen, T. O., Winfield, I. J., Vøllestad, L. A., Fletcher, J. M., James, J. B., & Stenseth, N. C. (2006). The ideal free pike: 50 years of fitness-maximizing dispersal in Windermere. Proceedings of the Royal Society B, 273, 2917–2924. doi:10.1098/rspb.2006.3659

Haugen, T. O., Winfield, I. J., Vøllestad, L. A., Fletcher, J. M., James, J. B., & Stenseth, N. C. (2007). Density dependence and density independence in the demography and dispersal of pike over four decades. Ecological Monographs, 77, 483–502. doi:10.1890/06-0163.1

Hunt, L. M. (2005). Recreational fishing site choice models: insights and future opportunities. Human Dimensions of Wildlife, 10, 153–172. doi:10.1080/10871200591003409

Hunt, L. M., Boxall, P., Englin, J., & Haider, W. (2005). Remote tourism and forest management: a spatial hedonic analysis. Ecological Economics, 53, 101–113. doi:10.1016/j.ecolecon.2004.06.025

Hunt, L. M., Arlinghaus, R., Lester, N., & Kishneriuk, R. (2011). The effects of regional angling effort, angler behavior, and harvesting efficiency on landscape patterns of overfishing. Ecological Applications, 21, 2555–2575. doi:10.1890/10-1237.1

Hunt, L. M., Sutton, S. G., & Arlinghaus, R. (2013). Illustrating the critical role of human dimensions research for understanding and managing recreational fisheries within a social-ecological system framework. Fisheries Management and Ecology, 20, 111–124. doi:10.1111/j.1365-2400.2012.00870.x

Hyman, A. A., McMullin, S. L., & DiCenzo, V. (2016). Dispelling assumptions about stocked-trout fisheries and angler satisfaction. North American Journal of Fisheries Management, 36, 1395–1404. doi:10.1080/02755947.2016.1221003

Johnson, B. M., & Carpenter, S. R. (1994). Functional and numerical responses: a framework for fish-angler interactions? Ecological Applications, 4, 808–821. doi:10.2307/1942010

Johnston, F. D., Arlinghaus, R., & Dieckmann, U. (2010). Diversity and complexity of angler behaviour drive socially optimal input and output regulations in a bioeconomic recreational-fisheries model. Canadian Journal of Fisheries and Aquatic Sciences, 67, 1507–1531. doi:10.1139/F10-046

Johnston, F. D., Arlinghaus, R., Stelfox, J., & Post, J. R. (2011). Decline in angler use despite increased catch rates: Anglers’response to the implementation of a total catch-and-release regulation. Fisheries Research, 110, 189–197. doi:10.1016/j.fishres.2011.04.006

Johnston, F. D., Arlinghaus, R., & Dieckmann, U. (2013). Fish life history, angler behaviour and optimal management of recreational fisheries. Fish and Fisheries, 14, 554–579. doi:10.1111/j.1467-2979.2012.00487.x

Johnston, F. D., Beardmore, B., & Arlinghaus, R. (2015). Optimal management of recreational fisheries in the presence of hooking mortality and noncompliance – predictions from a bioeconomic model incorporating a mechanistic model of angler behavior. Canadian Journal of Fisheries and Aquatic Sciences, 71, 37–53. doi:10.1139/cjfas-2013-0650

Kaufmann, S. D., Snucins, E., Gunn, J. M., & Selinger, W. (2009). Impacts of road access on lake trout *(Salvelinus namaycush)* populations: regional scale effects of overexploitation and the introduction of smallmouth bass (*Micropterus dolomieu*). Canadian Journal of Fisheries and Aquatic Sciences, 66, 212–223. doi:10.1139/F08-205

de Kerkhove, D. T.,. Minns, C. K., & Chu, C. (2015). Estimating fish exploitation and aquatic habitat loss across diffuse inland recreational fisheries. PLoS One, 10, e0121895. doi:10.1371/journal.pone.0121895. doi:10.1371/journal.pone.0121895

Lester, N. P., Marshall, T. R., Armstrong, K., Dunlop, W. I., & Ritchie, B. (2003). A broad-scale approach to management of Ontario’s recreational fisheries. North American Journal of Fisheries Management, 23, 1312–1328. doi:10.1577/M01-230AM

Lester, N. P., Shuter, B. J., & Abrams, P. A. (2004). Interpreting the von Bertalanffy model of somatic growth in fishes: the cost of reproduction. Proceedings of the Royal Society B: Biological Sciences, 271, 1625–1631. doi:10.1098/rspb.2004.2778

Lester, N. P., Shuter, B. J., & Venturelli, P. (2014). Life-history plasticity and sustainable exploitation: A theory of growth compensation applied to walleye management. Ecological Applications, 24, 38–54. doi:10.1890/12-2020.1

Levin, S., Xepapadeas, T., Crépin, A.-S., & Norberg, J. (2013). Social-ecological systems as complex adaptive systems: Modeling and policy implications. Environment and Development Economics, 18, 111–132. doi:10.1017/S1355770X12000460

Little, L. R., & McDonald, A. D. (2007). Simulations of agents in social networks harvesting a resource. Ecological Modelling, 204, 379–386. doi:10.1016/j.ecolmodel.2007.01.013

Mace, P. M. (1994). Relationships between common biological reference points used as thresholds and targets of fisheries management strategies. Canadian Journal of Fisheries and Aquatic Sciences, 51, 110–122. doi:10.1139/f94-013

Martin, D. R., Pracheil, B. M., DeBoer, J. A., Wilde, G. R., & Pope, K. L. (2012). Using the internet to understand angler behavior in the information age. Fisheries, 37, 458–463. doi:10.1080/03632415.2012.722875

Massey, D. M., Newbold, S. C., & Gentner, B. (2006). Valuing water quality changes using a bioeconomic model of a coastal recreational fishery. Journal of Environmental Economics and Management, 52, 482–500. doi:10.1016/j.jeem.2006.02.001

Matsumura, S., Arlinghaus, R., & Dieckmann, U. (2010). Foraging on spatially distributed resources with sub-optimal movement, imperfect information, and travelling costs: Departures from the ideal free distribution. Oikos, 119, 1469–1483. doi:10.1111/j.1600-0706.2010.18196.x

Matsumura, S., Arlinghaus, R., & Dieckmann, U. (2011). Assessing evolutionary consequences of size-selective recreational fishing on multiple life-history traits, with an application to northern pike (*Esox lucius*). Evolutionary Ecology, 25, 711–735. doi: 10.1007/s10682-010-9444-8

McFadden, D. (1973). Conditional logit analysis of qualitative choice behavior. In P. Zarembka (Ed.), Frontiers of econometrics (pp. 105–142). New York: Academic Press.

Mee, J. A., Post, J. R., Ward, H., Wilson, K., Newton, E., & Cantin, A. (2016). Interaction of ecological and angler processes: experimental stocking in an open access, spatially structured fishery. Ecological Applications, 26, 1693–1707. doi:10.1890/15-0879.1

Minns, C. K., Randall, R. G., Moore, J. E., & Cairns, V. W. (1996). A model simulating the impact of habitat supply limits on northern pike, *Esox lucius,* in Hamilton Harbour, Lake Ontario. Canadian Journal of Fisheries and Aquatic Sciences, 53(Suppl. 1), 20–34. doi:10.1139/f95-258

Muoneke, M. I., & Childress, W. M. (1994). Hooking mortality: A review for recreational fisheries. Reviews in Fisheries Science, 2, 123–156. doi:10.1080/10641269409388555

Neilson, W., & Wichman, B. (2014). Social networks and non-market valuations. Journal of Environmental Economics and Management, 67, 155–170. doi:10.1016/j.jeem.2013.11.005

Newbold, S. C., & Massey, D. M. (2010). Recreation demand estimation and valuation in spatially connected systems. Resource and Energy Economics, 32, 222–240. doi:10.1016/j.reseneeco.2009.11.014

Parkinson, E. A., Post, J. R., & Cox, C. P. (2004). Linking the dynamics of harvest effort to recruitment dynamics in a multistock, spatially structured fishery. Canadian Journal of Fisheries and Aquatic Sciences, 61, 1658–1670. doi:10.1139/f04-101

Patterson W. F., & Sullivan, M. G. (2003). Testing and refining the assumptions of put-and-take rainbow trout fisheries in Alberta. Human Dimensions of Wildlife, 18, 340–354. doi:10.1080/10871209.2013.809827

Paukert, C. P., Klammer, J.A., Pierce, R. B., & Simonson, T. D. (2001). An overview of northern pike regulations in North America. Fisheries, 26, 6–13. doi:10.1577/1548-8446(2001)026%3C0006:AOONPR%3E2.0.CO;2

Persson, L., de Roos, A. M., & Bertolo, A. (2004). Predicting shifts in dynamics of cannibalistic field populations using individual-based models. Proceedings of the Royal Society of London B: Biological Sciences, 271, 2489–2493. doi:10.1098/rspb.2004.2854

van Poorten, B. T., Arlinghaus, R., Daedlow, K., & Haertel-Borere, S. S. (2011). Social-ecological interactions, management panaceas, and the future of wild fish populations. Proceedings of the National Academy of Sciences of the United States of America, 108, 12554–12559. doi:10.1073/pnas.1013919108

Post, J. R., Parkinson, E. A. (2012). Temporal and spatial patterns of angler effort across lake districts and policy options to sustain recreational fisheries. Canadian Journal of Fisheries and Aquatic Sciences, 69, 321–329. doi:10.1139/f2011-163

Post, J. R., Sullivan, M., Cox, S., Lester, N. P., Walters, C. J., Parkinson, E. A., Paul, A. J., Jackson, L., & Shuter, B. J. (2002). Canada’s recreational fishery: The invisible collapse? Fisheries, 27, 6–17. doi: 10.1577/1548-8446(2002)027<0006:CRF>2.0.CO;2

Post, J. R., Mushens, C., Paul, A., & Sullivan, M. (2003). Assessment of alternative harvest regulations for sustaining recreational fisheries: model development and application to bull trout. North American Journal of Fisheries Management, 23, 22–34. doi:10.1577/1548-8675(2003)023<0022:AOAHRF>2.0.CO;2

Post, J. R., Persson, L., Parkinson, E. A., & van Kooten, T. (2008). Angler numerical response across landscapes and the collapse of freshwater fisheries. Ecological Applications, 18, 1038–1049. doi:10.1890/07-0465.1

Raat, A. J. P. (1988). Synopsis of biological data on the northern pike Esox lucius Linnaeus, 1758. FAO Fish. Synop. 30, Rev. 2. Rome: FAO.

Rogers, M. W., Allen, M. S., Brown, P., Hunt, T., Fulton, W., & Ingram, B. A. (2010). A simulation model to explore the relative value of stock enhancement versus harvest regulations for fishery sustainability. Ecological Modelling, 221, 919–926. doi:10.1016/j.ecolmodel.2009.12.016

Schlüter, M., McAllister, R. R. J., Arlinghaus, R., Bunnefeld, N., Eisenack, K., Hölker, F., Milner-Gulland, E. J., Müller, B., Nicholson, E., Quaas, M., & Stöven, M. (2012). New horizons for managing the environment: a review of coupled social-ecological systems modeling. Natural Resource Modeling, 1, 219–279. doi:10.1111/j.1939-7445.2011.00108.x

Shuter, B. J., Jones, M. L., Korver, R. M. & Lester, N. P. (1998). A general, life history based model for regional management of fish stocks: the inland lake trout *(Salvelinus namaycush)* of fisheries of Ontario. Canadian Journal of Fisheries and Aquatic Sciences, 55, 2161–2177. doi:10.1139/f98-055

Sullivan, M. G. (2002). Illegal angling of walleyes protected by length limits in Alberta. North American Journal of Fisheries Management, 22, 1053–1063. doi:10.1577/1548-8675(2002)022<1053:IAHOWP>2.0.CO;2

Swait, J. (1994). A structural equation model of latent segmentation and product choice for cross-sectional revealed preference choice data. Journal of Retail and Consumer Services, 1, 77–89. doi:10.1016/0969-6989(94)90002-7

Thébaud, O., Little, L. R., & Fulton, E. (2014). Evaluation of management strategies in Ningaloo Marine Park, Western Australia. International Journal of Sustainable Society, 6, 102–119. doi:10.1504/IJSSOC.2014.057892

Walters, C. J., & Martell, S. J. D. (2004). Fisheries ecology and management. Princeton, NJ: Princeton University Press.

Ward, H. G. M., Askey, P. J., & Post, J. R. (2013a). A mechanistic understanding of hyperstability in catch per unit effort and density-dependent catchability in a multistock recreational fishery. Canadian Journal of Fisheries and Aquatic Sciences, 70, 1542–1550. doi:10.1139/cjfas-2013-0264

Ward, H. G. M., Quinn, M. S., & Post, J. R. (2013b). Angler characteristics and management implications in a large, multistock, spatially structured recreational fishery. North American Journal of Fisheries Management, 33, 576–584. doi:10.1080/02755947.2013.785991

Ward, H. G. M., Allen, M. S., Camp, E. V., Cole, N., Hunt, L. M., Matthias, B., Post, J. R., Wilson, K., & Arlinghaus, R. (2016). Understanding and managing social–ecological feedbacks in spatially structured recreational fisheries: The overlooked behavioral dimension. Fisheries, 41, 524–535. doi:10.1080/03632415.2016.1207632

Wilberg, M. J., Irwin, B. J., Jones, M. L., & Bence, J. R. (2008). Effects of source–sink dynamics on harvest policy performance for yellow perch in southern Lake Michigan. Fisheries Research, 94, 282–289. doi:10.1016/j.fishres.2008.05.003

Wierzbicki, A. P. (1982). A mathematical basis for satisficing decision making. Mathematical Modelling, 3, 391–405. doi:10.1016/0270-0255(82)90038-0

Wilde, G. R., & Ditton, R. B. (1994). A management-oriented approach to understanding diversity among largemouth bass anglers. North American Journal of Fisheries Management, 14, 34–40. doi:10.1577/1548-8675(1994)014<0034:AMOATU>2.3.CO;2

Willis, D. W. (1989). Proposed standard length-weight equation for northern pike. North American Journal of Fisheries Management, 9, 203–208. doi:10.1577/1548-8675(1989)009<0203:PSLWEF>2.3.CO;2

Wilson, K. L., Cantin, A., Ward, H. G. M., Newton, E. R., Mee, J. A.,Varkey, D., Parkinson, E., & Post, J. R. (2016). Supply-demand equilibria and the size-number trade-off in spatially structured recreational fisheries. Ecological Applications, 26, 1086–1097. doi:10.1890/14-1771

Worm, B., Hilborn, R., Baum, J. K., et al. (2009). Rebuilding global fisheries. Science, 325, 578–585. doi:10.1126/science.1173146

Ziegler, J. P., Golebie, E. J., Jones, S. E., Weidel, B. C., & Solomon, C. T. (2017). Social-ecological outcomes in recreational fisheries: the interaction of lakeshore development and stocking. Ecological Applications, 27, 56–65. doi:10.1002/eap.1433

## References

Aas, Ø., & Ditton, R. B. (1998). Human dimensions perspective on recreational fisheries management: implications for Europe. In P. Hickley, & H. Tompkins (Eds.), Recreational fisheries. Social, economic and management aspects (pp. 153–164). Oxford, UK: Blackwell Publishing Ltd.

Anderson, D. K., Ditton, R. B., & Hunt, K. M. (2007). Measuring angler attitudes toward catch-related aspects of fishing. Human Dimensions of Wildlife, 12, 181–191.

Arlinghaus, R., & Mehner, T. (2003). Determinants of management preferences of urban anglers: Habitat rehabilitation versus other options. Fisheries, 28, 10–17.

Arlinghaus, R., Beardmore B., Riepe, C., Meyerhoff, J., & Pagel, T. (2014). Species-specific preferences of German recreational anglers for freshwater fishing experiences, with emphasis on the intrinsic utilities of fish stocking and wild fishes. Journal of Fish Biology, 85, 1843–1867.

Arterburn, J. E., Kirby, D. J., & Berry, C. R. (2002). A Survey of angler attitudes and biologist opinions regarding trophy catfish and their management. Fisheries, 27, 10–21.

Beardmore, B., Haider, W., Hunt, L. M., & Arlinghaus, R. (2013). Evaluating the ability of specialization indicators to explain fishing preferences. Leisure Science, 35, 273–292.

Bryan, H. (1977). Leisure value systems and recreational specialization: The case of trout fishermen. Journal of Leisure Research, 9, 174–187.

Buchanan, T. (1985). Commitment and leisure behavior: A theoretical perspective. Leisure Sciences, 7, 401–420.

Dorow, M., & Arlinghaus, R. (2011). A telephone-diary-mail approach to survey recreational fisheries on large geographic scales, with a note on annual landings estimates by anglers in northern Germany. In T. D. Beard Jr., R. Arlinghaus, & S. Sutton (Eds.) Proceedings from the fifth world recreational fishing conference. American Fisheries Society Symposium 75 (pp. 319–344). Bethesda, MD: American Fisheries Society.

Dorow, M., Beardmore, B., Haider, W., & Arlinghaus, R. (2010). Winners and losers of conservation policies for European eel, *Anguilla anguilla:* An economic welfare analysis for differently specialised eel anglers. Fisheries Management and Ecology, 17, 106–125.

Fedler, A. J., & Ditton, R. B. (1986). A framework for understanding the consumptive orientation of recreational fishermen. Environmental Management, 10, 221–227.

Graefe, A. R. (1980). The relationship between level of participation and selected aspects of specialization in recreational fishing. (Doctoral Dissertation). Texas A&M University, College Station.

McFadden, D. (1973). Conditional logit analysis of qualitative choice behavior. In P. Zarembka (Ed.) Frontiers of Econometrics (pp. 105–142). New York: Academic Press.

Oh, C., & Ditton, R. B. (2006). Using recreation specialization to understand multi-attribute management preferences. Leisure Sciences, 28, 369–384.

Salz, R. J., & Loomis, D. K., (2005). Recreation specialization and anglers’ attitudes towards restricted fishing areas. Human Dimensions of Wildlife, 10, 187–199.

Scherer, K. R. (2005). What are emotions? And how can they be measured? Social Science Information, 44, 695–729.

Scott, D., & Shafer, C. S. (2001). Recreational specialization: A critical look at the construct. Journal of Leisure Research, 33, 319–343.

Shafer, E. L. Jr. (1969). The average camper who doesn’t exist. Upper Darby, PA: U.S. Forest Service, Northeastern Forest Experiment Station.

